# scClassify: hierarchical classification of cells

**DOI:** 10.1101/776948

**Authors:** Yingxin Lin, Yue Cao, Hani J Kim, Agus Salim, Terence P. Speed, Dave Lin, Pengyi Yang, Jean Yee Hwa Yang

**Author notes:** Corresponding authors: Jean Y H Yang and Pengyi Yang.

## Abstract

Cell type identification is a key computational challenge in single-cell RNA-sequencing (scRNA-seq) data. To capitalize on the large collections of well-annotated scRNA-seq datasets, we present scClassify, a hierarchical classification framework based on ensemble learning. scClassify can identify cells from published scRNA-seq datasets more accurately and more finely than in the original publications. We also estimate the cell number needed for accurate classification anywhere in a cell type hierarchy.

---

Single cell RNA-sequencing (scRNA-seq) datasets have become larger and more diverse, driving the need for classification methods that can provide more refined discrimination between cell types. Many major cell types can be divided into subtypes in a hierarchical fashion, forming what we call a ‘cell type hierarchy’^1,2^. Approaches to scRNA-seq studies that account for cell type hierarchies in experimental design and cell type identification will permit more nuanced interpretations of the resulting data.^3^

The most common approach to identifying cell types within scRNA-seq data is unsupervised clustering^4–6^ followed by manual annotation using known marker genes. However, the number of clusters is rarely known in advance, and the annotation of clusters is highly dependent on prior knowledge of previously identified marker genes. This can bias the analysis towards better characterised cell types. An alternative approach is to use supervised learning (Supplementary Table 1). Here we can take advantage of the increasing number of scRNA-seq datasets with high quality annotations to reduce the bias associated with choosing marker genes. Such reference datasets can be used to develop a cell type prediction model, which may then be used to identify cell types in new datasets. Current supervised learning approaches typically use a single reference dataset, and if a cell type in a new dataset does not exist in the reference dataset, it will have a forced assignment or be combined with other types. Finally, current supervised learning methods use a ‘one pass’ classification approach, which does not take into account differences that may exist within a given cell type. The ability to define and effectively use cell type hierarchies is key to maximizing our understanding of cell types and their responses.

To address these challenges, we developed scClassify, a classification framework for robust cell type classification and discovery (Fig. 1a). scClassify first constructs a cell type tree from a reference dataset, allowing for multiple cell types at each non-terminal node. Next, scClassify uses a combination of gene selection methods and similarity metrics to develop classifiers that capture cell type characteristics at each such node. These classifiers are then integrated to make predictions for every cell at each node of the cell type tree. Depending on the numbers of cells in the reference dataset from each subtype at a branch node of the tree, scClassify may assign a cell from a new dataset to an intermediate or non-terminal cell type, and not classify it any further in the hierarchy. It also allows cells from a new dataset to be unassigned, reflecting the possibility that such cells may be of a type that is not in the reference dataset (Supplementary Fig. 1a). For these unassigned cells, scClassify uses a clustering procedure for novel cell type discovery (Fig. 1a). scClassify also enables classification of cells in a new dataset using multiple reference datasets (Fig. 1b), which can improve classification accuracy and reduce the number of intermediate and unassigned cells. Furthermore, scClassify addresses an important design challenge: estimating the number of cells required for accurately discriminating between cell subtypes anywhere in a cell type hierarchy.

**Fig. 1.**
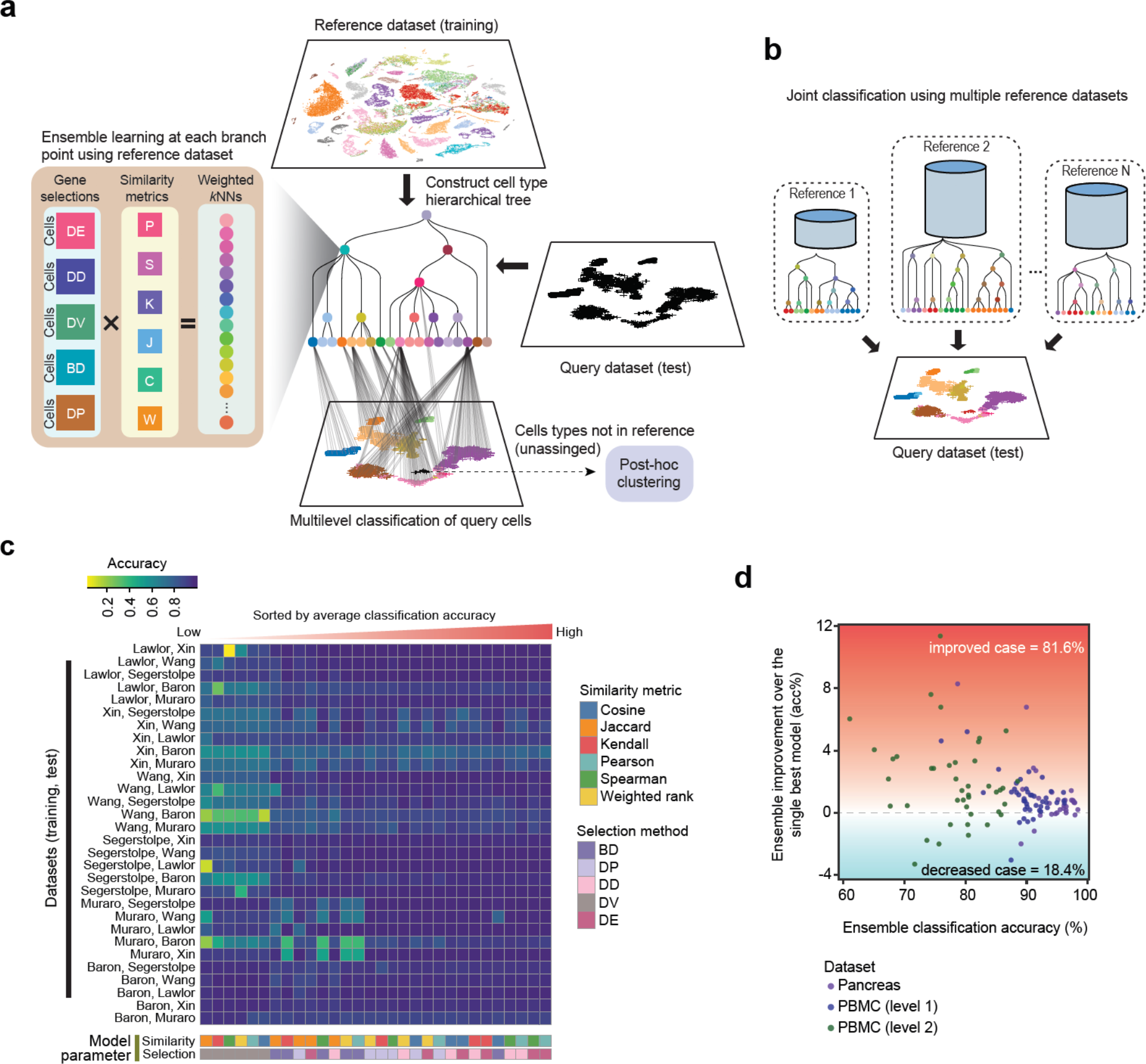
**a**, Schematic illustration of the scClassify framework. **b**, Schematic illustration of the joint classification using multiple reference datasets. **c**, Classification accuracy of all pairs of reference and test datasets was calculated using the all combinations of six similarity metrices, five gene selection methods. **d**, Improvement in classification accuracy after applying an ensemble learning model compared to the best single model (weighted kNN+Pearson+DE).

To assess the performance of scClassify, we collated seven PBMC datasets^7^ generated by different protocols and six publicly available human pancreas scRNA-seq datasets (Supplementary Fig. 1b). We first evaluated the performance of 30 individual classifiers on the pancreas data collection by training each on one dataset and testing it on another, with a weighted k-nearest neighbours (kNN) classifier using one of 5 gene selection methods and one of 6 similarity metrics (Fig. 1c). The heatmap highlights the diversity in performance across different parameter settings (with average accuracy ranging from 72% to 93%), suggesting that different parameter combinations capture different cell type characteristics and so the value of ensemble learning. While the differential expression (DE) gene selection method followed by weighted kNN with Pearson’s similarity metric is the best single classifier, the ensemble of weighted kNN classifiers trained by all 30 combinations of gene selection methods and similarity metrics led, in most cases, to a classification accuracy higher than that achieved by the single best model (Fig. 1d). The ensemble classifier is therefore used in all benchmarking experiments.

We compared the performance of scClassify with 11 other methods (Supplementary Table 1). The 6×5 = 30 (training, test) pairs from the pancreas data collection of 6 studies came in two groups: easy cases (n=16), where all cell types in the test data are found in the training data, and hard cases (n=14), where the test data contained one or more cell types not present in the training data. The results are summarised in Fig. 2a and Supplementary Figs. 2, 3. On average, scClassify achieves a higher accuracy than the other methods, with the difference being greater among the 14 hard cases than the 16 easy ones. For the collection of 6 PBMC datasets, we evaluated scClassify at two levels of the cell type hierarchy, coarse (“level 1”) or fine (“level 2”), each leading to 7×6=42 (training, test) pairs. We found that scClassify effectively annotates the cell types and produces higher accuracy rates in most cases (Fig. 2b and Supplementary Fig. 3), with the improvement being greater at level 2. Results for all methods are summarised in Supplementary Fig. 4. To test scClassify with a large dataset having a complex cell type hierarchy, we used the Tabula Muris FACS dataset^8^ as reference and the Microfluidic dataset (Fig. 2c, Supplementary Figs. 5, 6) as query, achieving a high concordance (~85% accuracy). Similar results were found (Supplementary Figs. 7, 8) for three mouse neuronal datasets^9,10^, each with about 20 cell types.

**Fig. 2.**
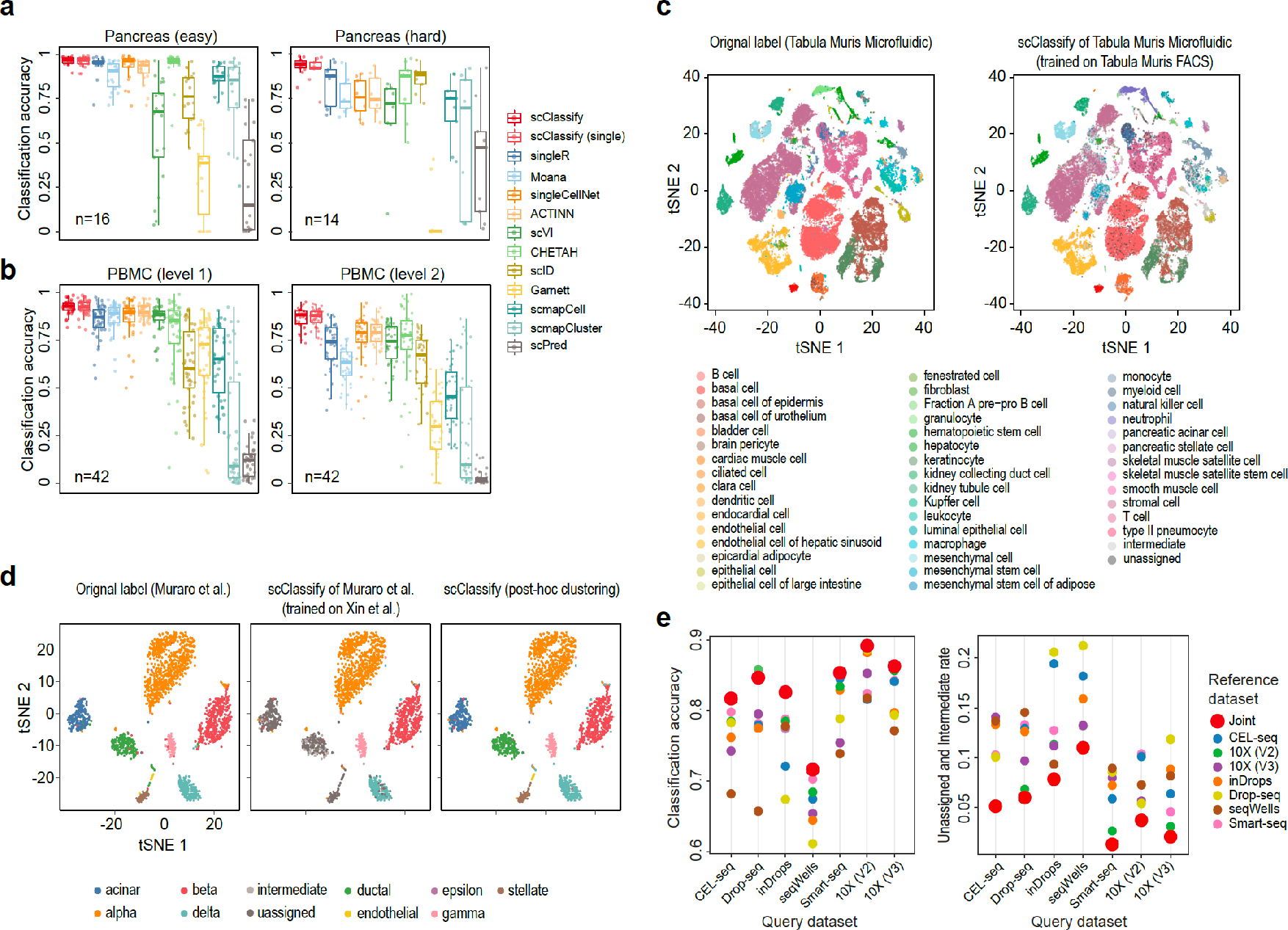
**a**, Performance evaluation for 13 methods on 30 training and test data pairs based on the Pancreas data collection. **b**, Performance evaluation for 13 methods on 84 training and test data pairs based on the PBMC data collection. **c**, tSNE visualisation of the Tabula Muris Microfluidic dataset. Cell typing is based on either the original publication (left panel; The Tabula Muris Consortium ^8^) or scClassify prediction (right panel). scClassify was applied to Tabula Muris Microfluidic data collection with Tabula Muris FACS dataset as training reference. **d**, Post-hoc cell typing of unassigned cells in scClassify predictions. Left panel shows cells types on the basis of the original publication by Muraro et al. ^20^. Middle panel shows the predicted cell types using scClassify trained using the reference dataset by Xin et al. as reference ^12^. Note that the reference dataset does not contain the cell types acinar, ductal and stellate cell types. Right panel shows post hoc clustering and cell typing results for unassigned cells found using scClassify prediction. **e**, Classifying query datasets using the joint prediction from multiple reference datasets (red circle). Classification accuracy of the joint prediction is compared to those obtained from using single reference datasets (other colours).

scClassify labels cells from a query dataset “unassigned” when their type is not in the reference dataset. With the Xin-Muraro pair^7,8^, scClassify correctly identified (Fig. 2d) the four shared cell types (alpha, beta, delta, and gamma) and correctly labelled as “unassigned” cells that were only present in Muraro dataset (i.e. acinar, ductal, stellate, endothelial, and delta). scClassify then applied scClust^11^ to the “unassigned” cells (Fig. 2d, Supplementary Fig. 9) and obtained clustering and labelling results consistent with those provided in the Muraro dataset. Another example is the Xin-Wang^12,13^ (reference, query) pair, see Supplementary Fig. 10. When multiple reference datasets are provided, scClassify can perform joint classification, resulting in higher classification accuracy compared to using single reference datasets, and fewer unassigned and intermediate cells (Fig. 2e).

As a complete design and identification framework, scClassify is able to estimate the number of cells required in a reference dataset to discriminate accurately between two cell types at a given level in the cell type hierarchy. It does so by fitting an inverse power-law^14^ (see on-line methods). The procedure requires no assumptions on the distributions of the training dataset or the accuracy. As expected, accuracy will increase with increasing sample size, and converge to a maximum. To evaluate this approach, we first select a given accuracy and the corresponding sample size from the learning curve estimated from the pilot data (Fig. 3a red line, Supplementary Fig. 11a). Then we perform an in-silico experiment by randomly selecting that sized sample of cells from full reference dataset, and build a cell type prediction model. Finally, the model is validated on an independent set of cells, and the corresponding in-silico experimental accuracy calculated (Fig. 3a, blue line, Supplementary Fig. 11a). The learning curve we estimate (Fig. 3a, red line) by this approach exhibits strong agreement (cor = 0.99) with the validation results, demonstrating the validity of the approach (Fig. 3b, Supplementary Fig. 11b).

**Fig. 3.**
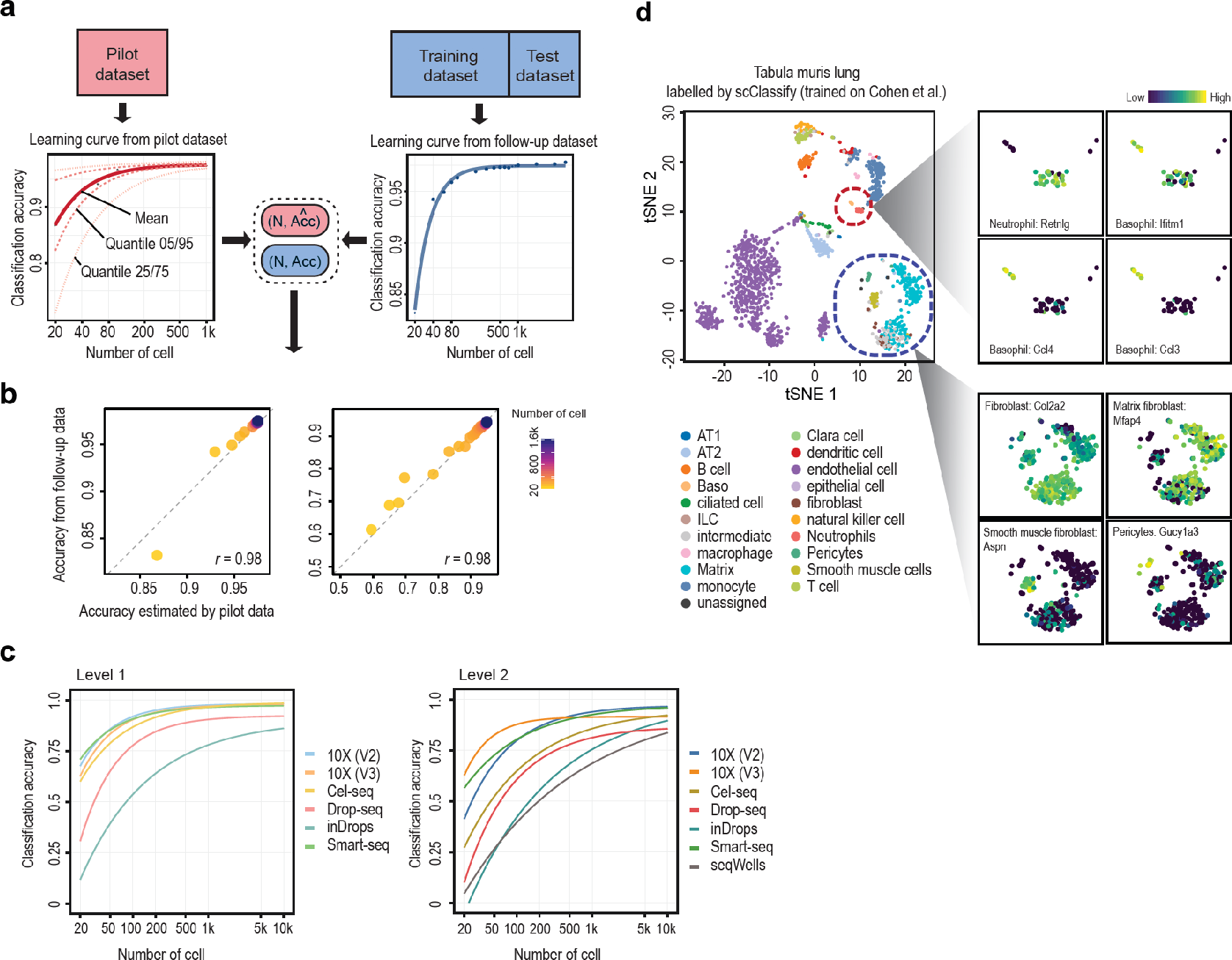
**a**, Schematic illustration of the scClassify sample size learning framework. **b**, Scatter plot of sample size estimation based on pilot data (horizontal axis) compared with accuracy results from in-silico experiments (vertical axis) **c**, Sample size learning curve with the horizontal axis representing sample size (N) and the vertical axis representing classification accuracy. The learning curves for the different datasets provide estimates of the sample size required to identify cell types at the top (left panel) and second (right panel) levels of the cell type hierarchical tree. **d**, scClassify results of the Tabula Muris Lung FACS data obtained using Cohen et al ^16^ dataset as reference. The large tSNE plot is the full data colored by scClassify-predicted cell types, while the smaller tSNE plots show two subsets of cells from the large plot, each coloured to highlight 4 different marker genes, where the lighter yellow color represents higher gene expression.

We applied the learning curve construction to PBMC datasets^7^ with different protocols and observed that most show very similar performance at the first level of the cell type hierarchy, with differences between protocols increasing at the second level of the tree (Fig. 3c). Moreover, through simulation using symSIM^15^ we find that within-population heterogeneity (sigma) and capture efficiency have a substantial impact on the sample size determination (Supplementary Fig. 13), while sequencing depth has little influence.

Finally, we illustrate the potential of scClassify to provide finely-grained annotation of cell types by refining the Tabula Muris Lung Atlas using as reference dataset an scRNA-seq study^15^ of mouse lung development with 20,931 cells across 22 cell types. scClassify showed that stromal cell in the original atlas can be sub-classified into smooth muscle and matrix fibroblast cells. Fig. 3d uses the key markers of each cell type reported by Cohen et al^16^ to validate our results, fibroblasts being marked by a high expression level of COL1A2; smooth muscle cells being marked by Aspn^17^, while matrix fibroblasts are marked by Macf2 and Mfap4. Furthermore, scClassify is able to meet the computational challenge of identifying the presence of small subpopulations of cells in an scRNA-seq dataset. For example, it identified a group of 6 basophil cells^18^ (marked by Mcpt8, Ccl3, Ccl4, and Ifitm1) originally labelled as leukocytes and dendritic cells. This ability to refine an annotation is again illustrated using a Tabula Muris kidney dataset as query and a different mouse kidney cell atlas data^19^ as reference (Supplementary Fig. 15, 16).

Summarizing, scClassify provides accurate and robust hierarchal cell type identifications setting using ensemble learning and post-hoc clustering. While there are several supervised learning methods for cell type identification in scRNA-seq datasets, scClassify has higher accuracy than these, and can refine published cell type identifications. Furthermore, scClassify provides a way to estimate the number of cells required for accurate cell type classification anywhere in a cell type hierarchy. The impact of scClassify is enhanced using multiple reference datasets, that can refine existing and subsequently annotate novel cell types.

## Methods

Methods, including statements of data availability and any associated accession codes and references, are available in the online version of the paper. The scClassify R package is available at https://sydneybiox.github.io/scClassify and the corresponding shiny app is at http://shiny.maths.usyd.edu.au/scClassify/.

## Author contributions

JYHY and PY conceived the study with input from YL. YL led the method development and data analysis with input from TPS, AS, PY and JYHY. YC and DL test and evaluate the method with input from HK. JYHY, PY, YL and YC interpreted the results with input from DL, TPS, AS and HK. YL implemented the R package with input from YC and HK. YC implemented the Shiny app with input from JYHY and YL. JYHY, PY, TPS and YL wrote the manuscript with input from all authors. All authors read and approved the final version of the manuscript.

## Acknowledgements

The authors thank all their colleagues, particularly at The University of Sydney, School of Mathematics and Statistics, for their support and intellectual engagement. We also thank Andy Tran for testing the package. The following sources of funding for each author, and for the manuscript preparation, are gratefully acknowledged: Australian Research Council Discovery Project grant (DP170100654) to JYHY, PY; Discovery Early Career Researcher Award (DE170100759) to PY; Australia NHMRC Career Developmental Fellowship (APP1111338) to JYHY. Research Training Program Tuition Fee Offset and Stipend Scholarship to YL, HK and YC. Chen Family Research Scholarship to YL. NIH grant (R21DC015107) to DL. The funding source had no role in the study design; in the collection, analysis, and interpretation of data, in the writing of the manuscript, and in the decision to submit the manuscript for publication.

## Methods

### scClassify framework

scClassify is a classification framework for identifying the cell types of cells in scRNA-seq studies. It uses a *cell type hierarchy* constructed from a reference dataset, and ensemble learning models at each *branch node* of the hierarchy trained on the reference dataset (Fig. 1a). Specifically, the hierarchical ordered partitioning and collapsing hybrid (HOPACH) algorithm [17] is used to construct the hierarchical cell type tree from the reference dataset (see *Cell type tree* section). In contrast to standard hierarchical clustering, this method allows a parent node to be partitioned into multiple child nodes, which is more consistent with the natural progression from broad to more specific cell types, where a cell type can have two or more subtypes.

We will use the term *training set* interchangeably with *reference set* (or *sets*), and restrict usage of the term test set to situations in which the true types of the cells whose type is being predicted are known, i.e. to situations where we are assessing or comparing the performance of scClassify. We will use the term *query set* or *query cell* when the unknown types of cells are to be predicted, and we write *test/query* when we wish to cover both cases.

scClassify uses a weighted k-nearest neighbour (kNN) classifier. More weight is given to neighbouring cells that are nearer, as defined by a similarity metric (see *Weighted kNN* section), to a query cell. To incorporate a variety of information, 6 similarity metrics and 5 cell type-specific gene selection methods are used in scClassify (see *Ensemble of base classifiers* section). Each of 30 base classifiers is trained using one of the similarity metrics and one of the gene selection methods. The final predictions are made by an ensemble classifier that weights individual classifiers depending on their training error. The best base classifier will have the highest weight, while a classifier with less than 50% accuracy will have negative weight.

An ensemble classifier is trained at each branch node of the cell type tree constructed using HOPACH, where both the training and test/query phase follow a top-down approach. That is, for parent nodes, classifiers are trained using the cells belonging to their child nodes. To characterise nodes at different levels, we carry out the gene selection step separately at each level of the tree. In the test/query step, a cell will have its type predicted from the highest to the lowest level, as long as it is possible to make a satisfactory prediction at every level (see *Multilevel classification section*). If that is not possible, the cell will be given an intermediate type. Finally, we carry out post-hoc clustering of unassigned cells.

#### Component 1: Cell type tree

To construct the cell type tree from a reference dataset, we first take the union across cell types of all the sets of genes found using *limma* [31] to be differentially expressed between one cell type and all other cell types (one-vs-all). We next use HOPACH on the average expression of selected genes from each cell type to construct the cell type tree. Starting from the tree *root*, the maximum number of children at each *branch node* was set as 5 by default, and can be modified when data containing a large number of cell types are used as reference. The root of the tree consists cells of all the cell types; the branch nodes of the tree are the less refined cell types, as each may contain multiple subtypes; at the bottom level of the tree are the *leaves*, representing the finest cell types in the reference dataset.

#### Component 2: Ensemble of base classifiers

At each branch node, an ensemble classifier is built from 30 base classifiers. Each is a weighted kNN model with a different combination of similarity metric and gene selection method (Fig. 1a). These 30 base classifiers are all combinations of six similarity metrics and five cell type-specific gene selection methods. The motivation is that while numerous similarity metrics are available, they often lead to different results reflecting the different properties of the data each metric is measuring [15, 34]. We use Pearson’s correlation, Spearman’s correlation, Kendall’s rank correlation, cosine distance, Jaccard distance, and weighted ranked correlation [13]. Since not all genes are cell type-specific [20], only the ones that are informative for cell type classification should be included in a similarity metric. There are various computational methods for selecting genes that are cell type-specific [35]. Five of them are included in scClassify for the different types of information each of them provides, four being based on two-sample tests. These are differentially expressed (DE) genes using *limma* [31] differentially variable (DV) genes using Bartlett’s test, differentially distributed (DD) genes using the Kolmogorov–Smirnov test, bimodally distributed (BD) genes using the bimodality index [41], and genes with differentially expressed proportions (DP) using a chi-squared test. To make a consensus prediction from an ensemble, we weight each base classifier in a manner similar to that in AdaBoost [4]. Specifically, the weight of each classifier is calculated as:

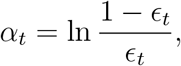

where *∊*_*t*_ is the error rate achieved by training and testing the base classifier *t* on the reference dataset, *t* = 1, …, 30. Let 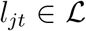 be the cell type label predicted for cell *j* by base classifier *t*, where 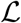 is the set of all the possible labels available to scClassify, including intermediate mode labels and ‘unassigned’. The ensemble prediction for cell *j* is made as [4]:

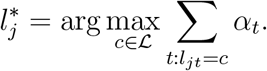

##### Weighted kNN

To relate the cell type predicted for a test/query cell to that of its nearest neighbours in the reference dataset, we use a distance-weighted kNN classifier [10].

Let 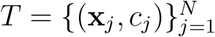 denote the reference data for a branch node, where 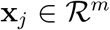 is the expression vector of cell-type specific genes for cell *j*, and *c*_*j*_ is the corresponding cell type label. For a test/query cell with expression vector 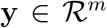, we first calculate the distances between it and all cells in the training dataset. Let *d*_*k*_ be the distance between such a cell and its *k*-th nearest neighbour in the reference dataset, *k* = 1, …, *N* (See *Similarity metrics* section). We then identify the *K* nearest cells in the training dataset for this cell. A weight *w*_*j*_ attributed to the *k*-th nearest neighbour is defined as

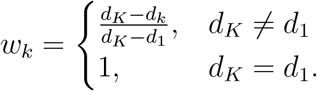

The query cell is then predicted to have the cell type with greatest total weight, i.e. we use weighted majority voting.

##### Similarity metrics

For simplicity in what follows, we identify cells with their (cell-type specific) gene expression vectors. Let 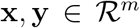 be cells with m gene expression values selected from the reference and query/test datasets, respectively. We consider the following metrics to measure dissimilarity between these two cells:

1. Pearson correlation

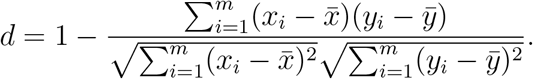
2. Spearman correlation

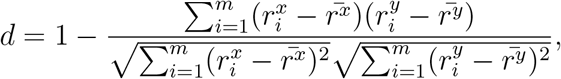

where 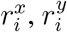 denote the rank of the expression value of gene *i* in cell **x**, **y** respectively; and *r̅* indicates the the mean rank of expression of the cell.
3. Kendall rank correlation

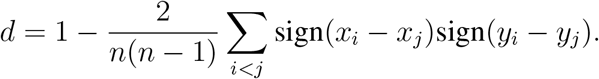
4. Cosine distance

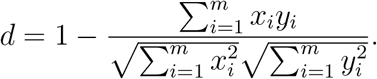
5. Jaccard distance

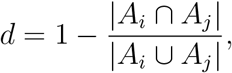

where *A*_*i*_, *A*_*j*_ indicate the set of genes that with expression greater than zero in cell *i* and cell *j*.
6. Weighted ranked correlation Weighted ranked correlation is simply Pearson’s correlation between the Savage scores of two cells [13]. This is calculated as follows:

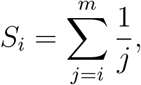

where *i* is the rank assigned to the *i*-th largest of the *m* gene expression values. Here, we are giving higher weight to agreement on the top rankings.

Note that the cosine and Jaccard distances are calculated using the *proxy* package [26]. scClassify uses Pearson’s correlation by default.

##### Discriminative genes

To characterise branch nodes in the tree, we select genes at each node separately. In all the methods below we compare the gene expression levels in each cell type at a node with those of all other cell types at that node, and then take the union across cell types of all the genes so selected. In 1-4 below, genes are then ranked according to their adjusted *p*-values for the test in question. In 1-3, we also require a gene’s PD as defined in 4 to exceed 0.05. We consider the following gene selection methods:

1. Differentially expressed (DE) genes: Differential expression analysis is carried out using limma-trend [31]. Using the *topTable* command of the *limma* package, we extract the genes with fold change > 1.
2. Differential variable (DV) genes: Differential variability analysis is carried out using Bartlett’s test.
3. Differentially distributed (DD) genes: Differential distribution analysis is carried out using the Kolmogorov–Smirnov test.
4. Differentially proportioned (DP) genes: Differential proportion analysis is performed by a chi-squared test. Here, each gene is classified as expressed or not expressed in each relevant cell, where expressed means it has an expression level greater than a certain threshold (by default, this threshold is set at 1 for log-transformed data). The difference between the proportion of cells in one cell type in which a gene is expressed and that proportion across all other cell types at that node is denoted by PD.
5. Bimodally distributed (BD) genes: We calculated the bimodality index [41] for each gene, as follows

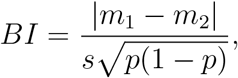

where *m*_1_, *m*_2_ are the means of the expression levels of one cell type and all the other cell types, *s* is the (assumed) common standard deviation, and *p* is the proportion of cells of the cell type under consideration. The genes are then ranked based on the bimodality index.

For each of these methods, we select a maximum of 50 top-ranked genes whose adjusted p-values are less than 0.01. These genes are included in training model.

#### Component 3: Multilevel classification

Starting from the root of the cell type tree, scClassify calculates the distances between a query cell and all reference cells at and below a branch node, and classifies the query cell to a child of that node if the following two criteria are fulfilled. First, the nearest neighbour cells must have correlations higher than a certain threshold. This is determined using a mixture model on the correlation of the cell type of this nearest neighbour, using the *normalmixEM* function in the package *mixtools* [5]. Second, the weights of its assigned cell type must be larger than a certain threshold, whose default is set at 0.7. Cells that fail to pass either of these criteria will not be classified to the next level. Cells that do not progress above the root will be considered *unassigned*. Those cells that are classified at a branch node, but whose classification process does not reach a leaf of the tree, will be viewed as having an *intermediate* cell type. In such cases, the final type of a query cell is defined by the last assigned branch node, where a branch node cell type is defined as the collection of cell types of all its child nodes. Finally, cells that reach the leaf level will be labelled by the cell type each leaf represents.

#### Component 4: Post-hoc clustering of unassigned cells

scClassify uses a modified version of the SIMLR algorithm [40] to cluster unassigned cells. To illustrate scClassify’s capacity to annotate cells that are not in the reference dataset, we trained scClassify on a reference dataset that had only four cell types (Xin et al.), and used this to predict cell types for a dataset with nine cell types from human pancreas (Muraro et al.). Cells that were classified as unassigned were then clustered and further identified in the familiar way following clustering. Specifically, DE genes in each cluster (one-vs-all) were found using *limma* [31]. Each cluster was subsequently annotated based on its DE genes and the markers provided by Muraro et al. in the paper (i.e. acinar: PRSS1; ductal: SPP1, KRT19; stellate: COL1A1, COL1A2; endothelial: ESAM; delta: SST) (Supplementary Fig. 9). Similarly, we performed post-hoc clustering on the human pancreas dataset generated by Wang et al., where most of ductal and stellate cells are correctly called unassigned by scClassify (Supplementary Fig. 10a). We annotated each cluster based on its genes DE when compared to other clusters (ductal: SPP1; stellate: COL1A2) (Supplementary Fig. 10b). In both cases, we found that the cell types assigned by scClassify were highly consistent with those originally published.

#### Component 5: Joint classification using different training datasets

When multiple scRNA-seq datasets are available, each profiling the same tissues or including overlapping cell types, scClassify can make use of them all. It will train a collection of ensemble models using each dataset and make predictions for a query dataset using a joint classification method. Predictions from each reference dataset are weighted by their training error, because classifiers with larger training errors usually make poorer predictions, and therefore should be down-weighted. For each query cell, scClassify assigns the cell type with the largest average score using the ensemble models from each of the reference datasets.

### Data collections and data processing

We organized a set of public datasets into large data collections for either performance assessment or use as case studies (Supplementary Fig. 1b). The pancreas data collection was downloaded from the National Center for Biotechnology Information (NCBI) Gene Expression Omnibus (GEO) for GSE81608 (Xin), GSE83139 (Wang), GSE86469 (Lawlor), GSE85241 (Muraro), GSE84133 (Baron), and EBI Array-Express website for E-MTAB-5061 (Segerstolpe) [43, 42, 43, 18, 28, 3, 33]. We manually checked the cell type annotations that were provided by the original authors of each dataset and curated the labels so that the naming conventions were consistent across datasets. For example, the cell type “PP” in Xin dataset was changed to “gamma”, as “gamma” was the name used by all other datasets. Similarly, “mesenchymal” in Muraro and Wang was changed to “stellate”. We also removed the cell types that were labelled as ‘co-expression’ and ‘unclassified endocrine’ in Segerstolpe dataset. We manually checked the cell type annotations that were provided by the original authors of each dataset and curated the labels such that the naming convention is consistent across datasets. For example, the cell type “PP” in Xin dataset was changed to “gamma”, as “gamma” was the name used by all other datasets. Similarly, “mesenchymal” in Muraro and Wang was changed to “stellate”. We also removed the cell types that were labelled as ‘co-expression’ and ‘unclassified endocrine’ in Segerstople dataset.

The PBMC data collection [9] was downloaded from https://portals.broadinstitute.org/single_cell/study/SCP424/single-cell-comparison-pbmc-data. It contains a collection of seven datasets that were sequenced using different platforms (Smart-seq, Cel-seq, inDrops, dropSeqs, seqWells, 10X Genomics (V3), 10X Genomics (V2))). The Tabula Muris mouse data [32] was downloaded from https://tabula-muris.ds.czbiohub.org/. The neuronal data collection was downloaded from GEO accession number GSE71585 (Tasic (2016) [37]), GSE115746 (Tasic (2018) [38]), and GSE102827 (Hrvatin [12]). The mouse lung development dataset was downloaded from GEO accession number GSE119228 [7]. The healthy mouse kidney dataset was downloaded from GEO accession number GSE107585 [29]. The PBMC10k data collection generated by Cell Ranger version 3.0.0 was downloaded https://support.10xgenomics.com/single-cell-gene-expression/datasets/3.0.0/pbmc_10k_v3.

For all datasets described above, only cells that passed the quality control of the original publication and assigned cell types were included. We performed size factor standardization to the raw count matrices for each batch/dataset using the *normalize* function in the R package *scater* [24] and used the log-transformed gene expression matrices as inputs to scClassify and all other methods. For the PBMC10K data, we also removed the doublets from the dataset using *DoubletFinder* [25]. We labelled the cells following the approach proposed by 10X Genomics for PBMCs [46], which allowed us to annotate 11 PBMC cell types. In sample size calculation study, we considered “B”, “Monocyte”, “T”, “NK”, “CD34+” cell type as first level coarse annotation and then expanded “T” to “CD4+T” and “CD8+T” for second level finer annotation.

### Performance evaluation

We extend the evaluation framework introduced in [14] by describing the results of our predictions as “correctly classified”, “misclassified”, “intermediate” (correct and incorrect), “incorrectly unassigned”, “incorrectly assigned”, and “correctly unassigned” (Supplementary Fig. 1a).

For cells of a given cell type from a query dataset, we first consider whether this cell type is present in the reference dataset. If it is, we then consider whether these cells are classified to the leaf level in the cell type hierarchy. Cells that are classified to the leaf level are “correctly classified” if their predicted cell types match their annotated cell types in the original study; otherwise they are “misclassified”. Of cells not classified to the leaf level, we consider the unassigned to be “incorrectly unassigned”, while if they are assigned “intermediate” cell types, we check whether their “intermediate” types contain the annotated cell types from the original study. Those that are on the correct branch of the cell type tree are “correct intermediate”; otherwise they are considered to be “incorrect intermediate”.

Cells from a cell type that in the query dataset that is not present in the reference dataset can be either “incorrectly assigned” to a cell type in the reference dataset, or “correctly unassigned”.

#### Evaluation and comparison methods

To evaluate and compare the performance of scClassify, we obtained 11 other publicly available scRNA-seq classification methods (Supplementary Table 1). These packages were installed either through their official CRAN or Bioconductor website, where available, or from their GitHub page. Two collections of datasets, pancreas and PBMC, were used (Supplementary Fig. 1b).

For the pancreas datasets, we defined the *hard* cases to be the 14 (training, test) = (reference, query) set pairs, where the reference/training dataset has *fewer* cell types than the query/test dataset. The remaining 16 pairs were called *easy*. The evaluation of scClassify on the PBMC datasets was carried out at two levels of the cell type hierarchy, coarse (“level 1”) or fine (“level 2”). Each led to 42 (training, test) set pairs. For level 1, we combined CD16+ and CD14+ monocytes into the cell type of monocytes, and we combined CD8+ and CD4+ T cells into the cell type of T cells. For the level 2, we used the original cell types.

The evaluation procedure was as follows. For both the pancreas and the PBMC data collections, we used one of the datasets as reference or training set, and all the other datasets as query or test sets, on which the accuracy of cell type predictions could be calculated. This gives us 6 × 5 = 30 distinct (training, test) set pairs of pancreas datasets, and 7 × 6 = 42 distinct pairs of PBMC datasets for each of the two levels. For each of the 11 methods, we evaluated the performance of a total of 114 (training, test) set pairs. We used the default settings given in the package README or vignette for training each method. The Garnett method requires the specification of marker genes for model training. In this case, we used genes obtained from the published literature. The log-transformed size factor normalized gene expression matrix was used as the input for all models.

The results were the types predicted for all cells in the test/query datasets. To calculate classification accuracy, we compared the predicted cell types with those provided in the reference dataset. Methods that do not allow unassigned or intermediate prediction were assessed based on cells that are “correctly classified” and “misclassified” (Supplementary Fig. 1a). For methods that do allow unassigned predictions, “incorrectly unassigned”, “incorrectly assigned”, and “correctly unassigned” were also included for calculating classification accuracy. Finally, for methods that allows both unassigned and intermediate predictions, we considered “correct intermediate” as correct predictions in calculating classification accuracy. We also calculated a conservative classification accuracy by treating all “intermediate” predictions as incorrect (Supplementary Fig. 3).

### Case studies: Identification of small cell subpopulation

We refined the annotation of the Tabula Muris lung FACS data as a query set using a mouse lung development dataset from of E12.5 to Day7 having 20,931 cells [7] from 22 cell types as reference. scClassify correctly annotated the common cell types such as endothelial, natural killer, T and B cells. It also identified two subtypes of fibroblasts (stromal cells): smooth muscle cells and matrix fibroblasts. In order to do so, we used the key markers of these cell types reported by Cohen et al. to confirm our classifications [7]. Cells that are classified as fibroblasts have high expression levels of the key marker Col1a2. Those classified as smooth muscle cells also have Aspn highly expressed, while matrix fibroblasts exhibit high expression of Macf2 and Mfap4 (Fig. 3d). scClassify was also able to annotate three minor populations from these data: pericytes (19 cells, originally labelled as stromal), ciliated cells (21 cells) and neutrophils (28 cells, originally labelled leukocytes), these classifications being supported by the high expression levels of marker genes in these cell types (pericytes: Gucy1a3; neutrophils: Retnlg) (Fig. 3d). Remarkably, scClassify was able to identify a group consisting of only 6 cells from the data, originally labelled as leukocyte and dendritic cells, but expressing high levels of the three top basophil markers highlighted by Cohen et al. (Ccl3, Ccl4, and Ifitm1) (Fig. 3d). This illustrates the ability of scClassify to identify cell types present in very small numbers in the data, something which is usually extremely challenging to do with single cell data using only clustering.

Further, we used a healthy mouse kidney dataset [29] containing 14 main cell types as a reference dataset to annotate the both the SMART-seq and Drop-seq kidney data from the Tabula Muris study. scClassify identified several large groups of cells as ascending loop of Henle (LOH), distal convoluted tubule (DCT), and proximal tubule (PT) cells from both the SMART-seq and the Drop-seq data, supported by the corresponding high marker expression level (Supplementary Figs. 15-16). Furthermore, scClassify identified two subgroups of collecting duct cells: collecting duct principal cells (CD-PC) and collecting duct intercalated cells (CD-IC). It should be noted that the top two key markers of CD-IC, Atp6v1g3 and Atp6v0d2, were only present in 3 cells and 1 cell in FACS data respectively (Supplementary Fig. 15). Lastly, leukocytes labelled by the original Tabula Muris paper could be identified as macrophage, B cells, T cells, and Neutrophils by scClassify (Supplementary Figs. 15-16).

### Framework for sample size determination

#### Learning curve construction

Estimating the number of cells required in a scRNA-seq study in order to discriminate between two given cell types for a given platform is essentially sample size determination for a high-dimensional classification problem. Our approach is to construct a learning curve [8, 27] based on a pilot study, and estimate the empirical classification accuracy as a function of the size of the reference dataset. For each dataset of size *N*, we randomly divide the data into a training set of size n and a test set of size *N* − *n*, independently *T* times (*T* = 20), with the resulting accuracy rates being denoted by 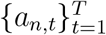. The average accuracy rate calculated by 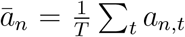. We then fit an inverse power-law function of sample size to the average accuracy rate as follows:

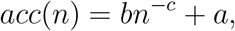

where *b* is the learning rate, *c* the decay rate, and *a* is the maximum accuracy rate that can be achieved by the classifier.

When we did this, we noticed that the inverse power-law function gave a poor fit in some cases where we had a small sample size. Therefore, we moved to a two component inverse power-law function to relate average accuracy rate to sample size, separately fitting to small and large sample sizes. That is, we now fit to

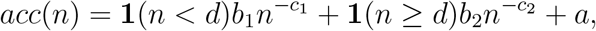

where *d* here is the transition point of the model, *b*_1_ and *b*_2_ are the learning rates of each component, and *c*_1_ and *c*_2_ their decay rates.

The function *nlsLM* in R package *minpack.lm* is used to fit both learning curves models [11]. To fit the mixture model, we first fixed *d* and estimated *b*_1_, *b*_2_, *c*_1_, *c*_1_, and *a*. After fitting the model for various values of *d*, we determined the best model to be the one with smallest residual sum of squares.

#### Learning curve construction with data integration

We used *scMerge* [21] to remove batch effects between two samples in the PBMC data collection generated from five different protocols [9] and next constructed the learning curves. We found that in most of the cases (SMART-seq, CEL-seq, and inDrops), the learning curve based on batch-corrected data achieved a higher accuracy rate that the uncorrected data (Supplementary Fig. 12), highlighting the need for batch effect removal within the training data.

### Evaluation of sample size determination for building the reference set

#### Evaluation of the learning curve

To validate the estimated learning curve, we randomly divide the PBMC10k data into the pilot (20%) and validation (80%) sets. The pilot set represents data we might have from a typical pilot study on which sample size estimation is based. The validation set represents data generated from an actual study. For this evaluation, the validation set is further divided into a training set and an independent test set, to calculate the accuracy of our learning curve for any given sample size. We hope and expect the learning curves constructed from these two subsets of the data will be consistent, and we will assess their consistency by the Pearson correlation between the accuracies estimated from the pilot data and those estimated from the validation data.

#### Sensitivity investigation of capture efficiency and sequencing depth via simulation

To investigate the potential influence of capture efficiency, sequencing depth and degree of cell type separation on sample size requirement, we carried out simulations using *SymSim* [45], estimating the parameters on PBMC10k data, with following parameter settings:

1. within-population heterogeneity(*σ*) from 0.2 to 1 in increments of 0.2. According to the definition by *SymSim*, a higher *σ* indicates a more homogeneous simulated population;
2. Capture efficiency at the following values 0.001, 0.01, 0.02, 0.03, 0.04, 0.05, 0.06, 0.07, 0.08, 0.09 and 0.1;
3. sequencing depth at the following values 30,000, 80,000, 160,000, 300,000 and 500,000.

#### Sensitivity investigation of capture efficiency and sequencing depth via downsampling

We first fitted the DECENT [44] model on the UMI data matrix for the PBMC10k data collection. After obtaining the parameter estimate of DECENT’s beta-binomial capture model, we conducted a UMI down-sampling using random draws from a beta-binomial distribution. Specifically, for the UMI raw count *x*_*ij*_ of gene *i* and cell *j*, the down-sampled matrix *Z*^(*k*)^ for down-sampling parameters *p*_*k*_ is generated by

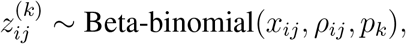

where *ρ*_*ij*_ is the correlation parameter of beta-binomial distribution, calculated using

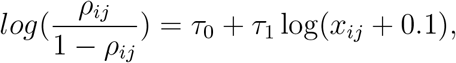

where *τ*_0_ and *τ*_1_ estimates of DECENT’s of the capture model parameters. Within each cell, the parameter *p*_*k*_ in this case can be interpreted as the ratio of capture efficiency in the down-sampled dataset relative to the original dataset.

We carried out 5-fold cross validation 20 times with each down-sampling proportion parameter, which ranges from 0.1 to 1 on PBMC10k data with two cell type levels. We found that for predictions at the top of cell type tree, scClassify achieved over 90% accuracy, even with 10% of original capture efficiency. For prediction at the second level of the cell type hierarchy, it requires 50% of original capture efficiency to achieve a similar level of performance (Supplementary Fig. 14a). We then constructed the learning curves of down-sampling matrices for *p*_*k*_ = 0.2, 0.5, 0.8, 1. As shown in Supplementary Fig. 14b, for level 1 prediction, learning curves converges to the highest accuracy rate at *N* = 80 when *p*_*k*_ is greater or equal to 0.5. However for level 2 prediction, it requires *N* = 400, and for the cases with 20% capture efficiency, the accuracy converges to about 75%.

## Supplementary Tables

**Table S1:**
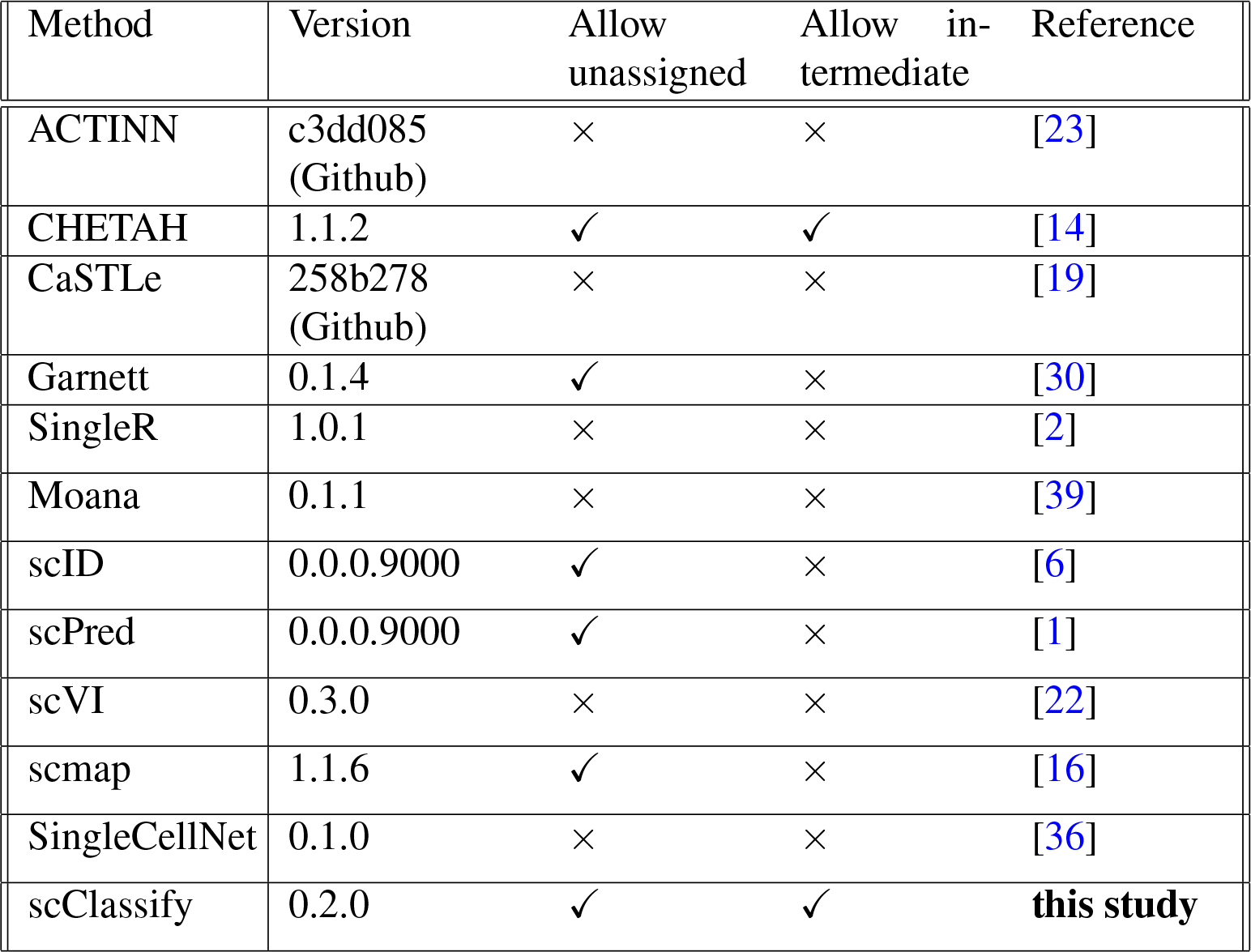
Current supervised learning methods for cell-type identification

