## Supplemental Figures for "scClassify: hierarchical classification of cells"

**a**

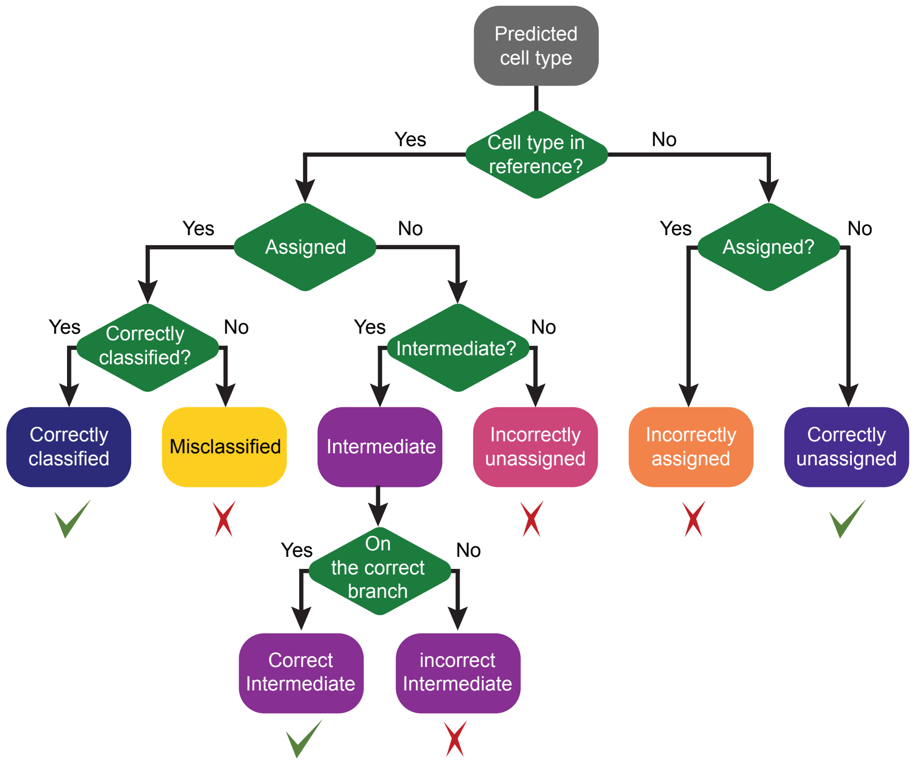


**b**

| | Data collection | Accession | Name | Protocol | Organism | Tissue | # of cell types | # of cells |
|---|---|---|---|---|---|---|---|---|
| Benchmark data collections | Pancreas | GSE81608<br>GSE83139<br>GSE86469<br>E-MTAB-5061<br>GSE85241<br>GSE84133 | Xin<br>Wang<br>Lawlor<br>Segerstolpe<br>Muraro<br>Baron | SMARTer/C1<br>SMART-seq<br>SMARTer/C1<br>SMART-seq2<br>Cel-seq2<br>inDrop | Human | Pancreas<br>Pancreas Islets | 4<br>7<br>7<br>11<br>9<br>13 | 1474<br>501<br>617<br>2127<br>2122<br>8569 |
| | PBMC | NA | Smart-seq<br>CEL-seq<br>10x (V2)<br>10x (V3)<br>inDrop<br>seqWells<br>Drop-seq | Smart-seq<br>CEL-seq<br>10x (V2)<br>10x (V3)<br>inDrop<br>seqWells<br>Drop-seq | Human | PBMC | 7<br>7<br>8<br>9<br>9<br>7<br>9 | 526<br>526<br>3362<br>3222<br>6584<br>6584<br>3727 |
| Case study data collections | Tabula Muris | GSE109774 | Tabula Muris FACS<br>Tabula Muris Microfluidic | FACS<br>Microfluidic | Mouse | Multiple | 68<br>40 | 41965<br>54439 |
| | Neuronal | GSE71585<br>GSE115746<br>GSE102827 | Tasic (2016)<br>Tasic (2018)<br>Hrvatin | SMARTer/C1<br>SMART-seq2<br>inDrop | Mouse | Primary visual cortex<br>Visual cortex | 20<br>23<br>20 | 1679<br>13586<br>48266 |
| | 10x PBMC | NA | 10x10k | 10x (V3) | Human | PBMC | 7 | 10753 |
| | Lung | GSE119228 | Cohen | MARS-seq | Mouse | Lung | 22 | 20931 |
| | Kidney | GSE107585 | Park | Drop-seq | Mouse | Kidney | 16 | 43745 |

Supplementary Figure 1: (a) Evaluation framework used in this study. Predictions are classified into "correctly classified", "misclassified", "intermediate" (either correct or incorrect), "incorrectly unassigned", "incorrectly assigned" or "correctly unassigned". (b) All datasets used in this study, including two collections for benchmarking and five collections for case study in sample size learning and rare cell type identification.

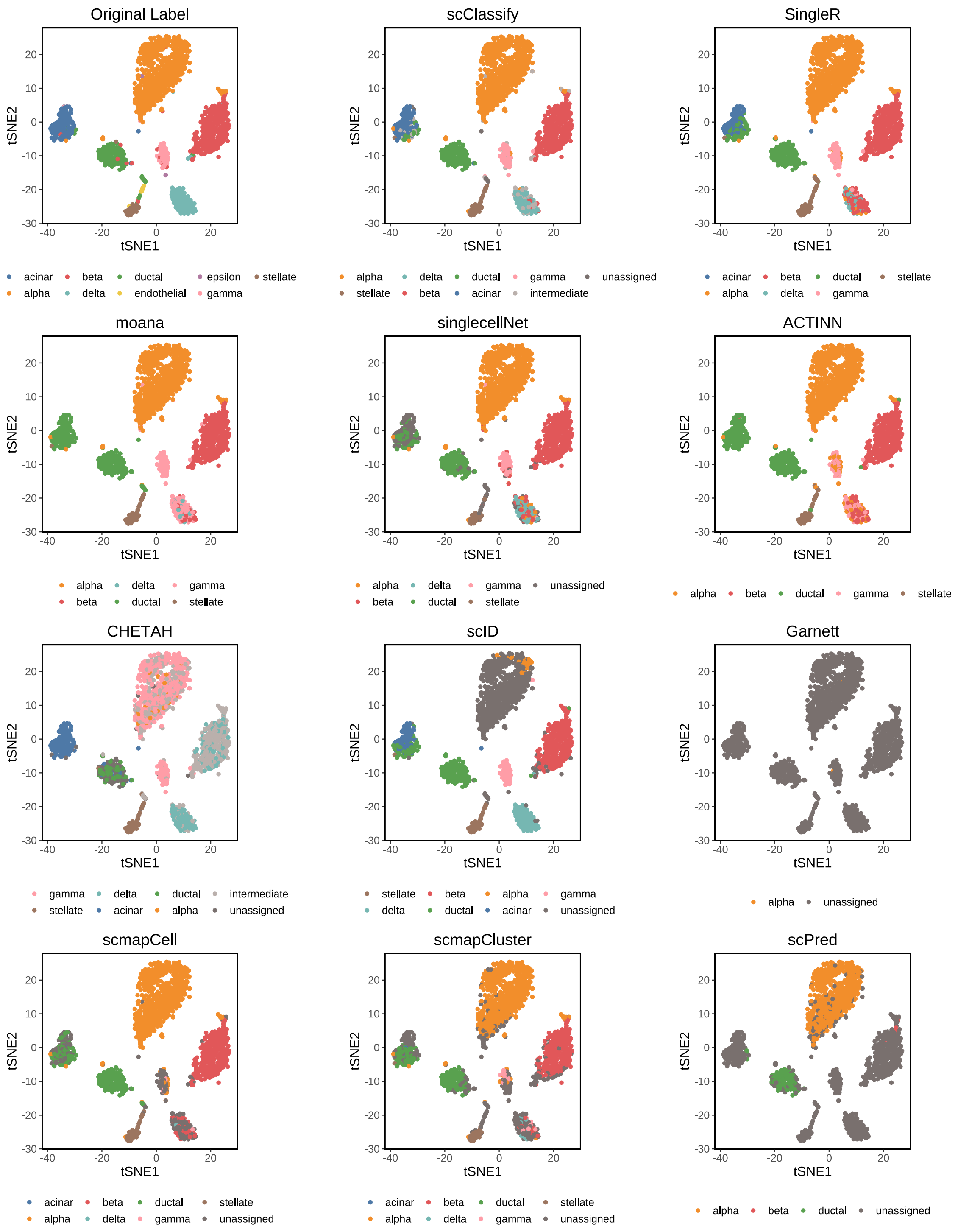


Supplementary Figure 2: A 4 by 3 panel tSNE plots of the Muraro et al. dataset from the human pancreas data collection. tSNE plots are color-coded by their original label (panel (1,1)) or predicted cell types from 11 different methods, scClassify, SingleR, moana, singlecellNet, ACTINN, CHETAH, scID, Garnett, scmap-cell, scmap-cluster, and scPred, which all used the Wang et al. dataset as the reference dataset (see Supp Table 1 for details). Under default settings, scClassify is able to correctly classify most cells with an accuracy rate of greater than 95%. None of the methods were fine-tuned.

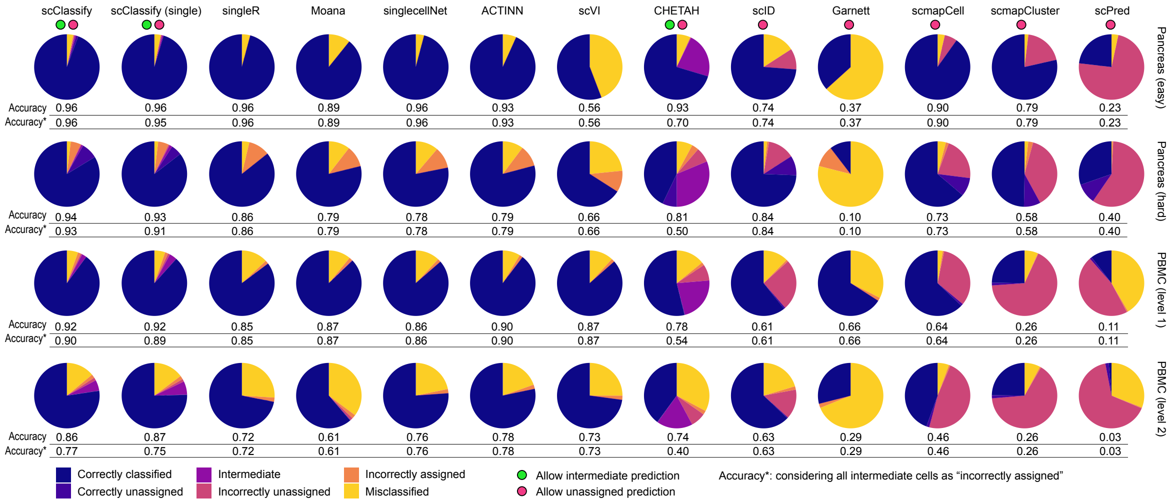


| | scClassify | scClassify (single) | singleR | Moana | singlecellNet | ACTINN | scVI | CHETAH | scID | Garnett | scmapCell | scmapCluster | scPred |
|---|---|---|---|---|---|---|---|---|---|---|---|---|---|
| Pancreas (easy) Accuracy | 0.96 | 0.96 | 0.96 | 0.89 | 0.96 | 0.93 | 0.56 | 0.93 | 0.74 | 0.37 | 0.90 | 0.79 | 0.23 |
| Pancreas (easy) Accuracy* | 0.96 | 0.95 | 0.96 | 0.89 | 0.96 | 0.93 | 0.56 | 0.70 | 0.74 | 0.37 | 0.90 | 0.79 | 0.23 |
| Pancreas (hard) Accuracy | 0.94 | 0.93 | 0.86 | 0.79 | 0.78 | 0.79 | 0.66 | 0.81 | 0.84 | 0.10 | 0.73 | 0.58 | 0.40 |
| Pancreas (hard) Accuracy* | 0.93 | 0.91 | 0.86 | 0.79 | 0.78 | 0.79 | 0.66 | 0.50 | 0.84 | 0.10 | 0.73 | 0.58 | 0.40 |
| PBMC (level 1) Accuracy | 0.92 | 0.92 | 0.85 | 0.87 | 0.86 | 0.90 | 0.87 | 0.78 | 0.61 | 0.66 | 0.64 | 0.26 | 0.11 |
| PBMC (level 1) Accuracy* | 0.90 | 0.89 | 0.85 | 0.87 | 0.86 | 0.90 | 0.87 | 0.54 | 0.61 | 0.66 | 0.64 | 0.26 | 0.11 |
| PBMC (level 2) Accuracy | 0.86 | 0.87 | 0.72 | 0.61 | 0.76 | 0.78 | 0.73 | 0.74 | 0.63 | 0.29 | 0.46 | 0.26 | 0.03 |
| PBMC (level 2) Accuracy* | 0.77 | 0.75 | 0.72 | 0.61 | 0.76 | 0.78 | 0.73 | 0.40 | 0.63 | 0.29 | 0.46 | 0.26 | 0.03 |

Supplementary Figure 3: Benchmarking results for 13 different methods. Each pie chart indicates the composition of predicted categories of the average performance in a collection of reference-testing pairs. We divided reference-test pairs into four groups: Pancreas (easy), Pancreas (hard), PBMC (level1), and PBMC (level2). Note that we only account for the intermediate cell types that have ended up in the right branch of the hierarchical tree. The accuracy rate is calculated as the sum of proportions of "correctly classified" and "correctly unassigned" and "intermediate on the correct path", while accuracy* indicates the sum of proportions of "correctly classified" and "correctly unassigned". Red and green dots indicate whether the methods allow cells to be classified as "intermediate" or "unassigned".

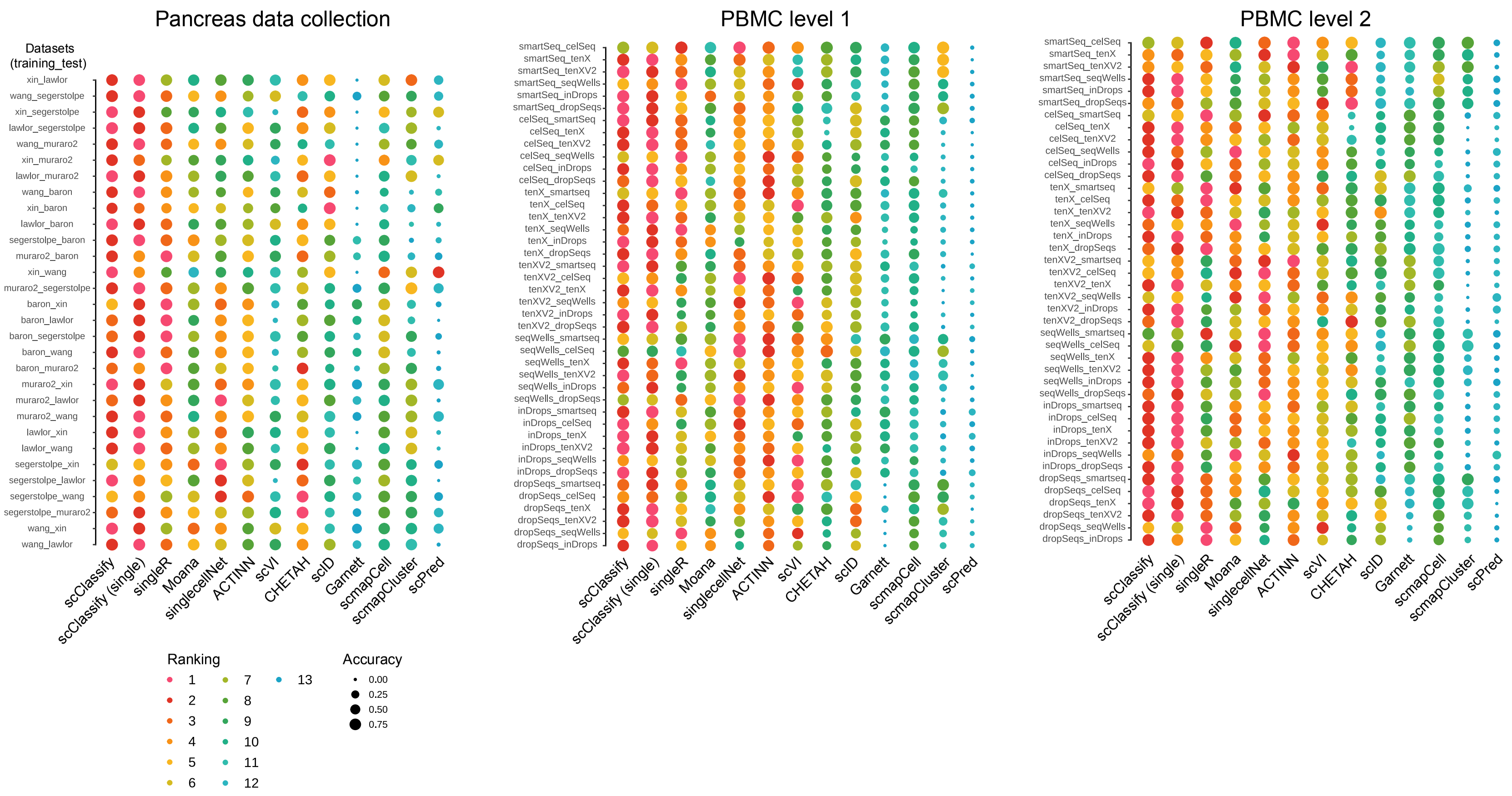


Supplementary Figure 4: A 1 by 3 panel of dot plots indicating the rankings of each method in 114 pairs of reference and testing data pairs. The x-axis refers to different methods and the y-axis refers to the reference and testing data pairs. The dots are colored by ranking and the size of the dots indicate the degree of accuracy.

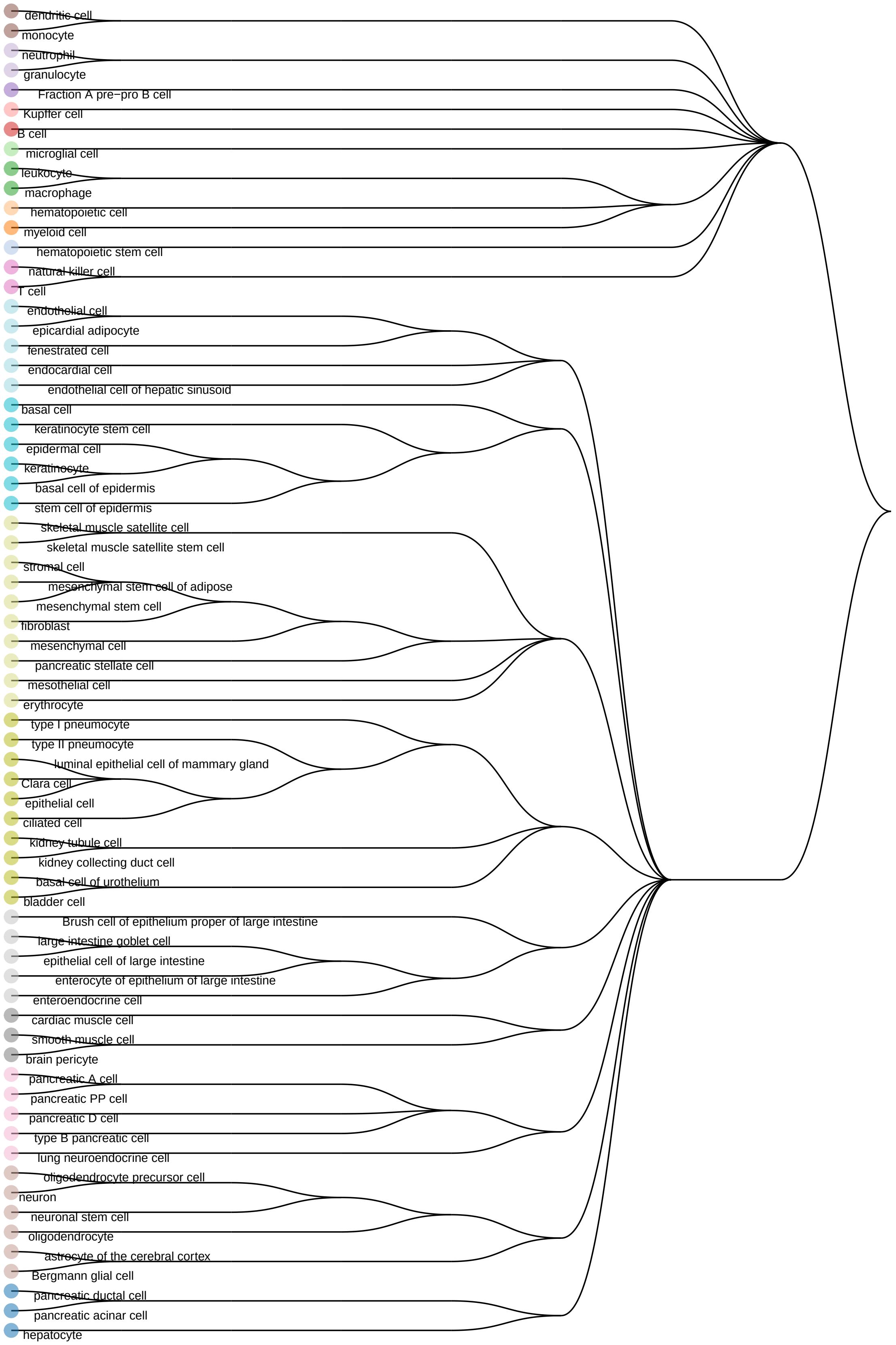


Supplementary Figure 5: A cell type tree generated using the hierarchical ordered partitioning and collapsing hybrid (HOPACH) algorithm [6] and the Tabula Muris FACS data as the reference dataset.

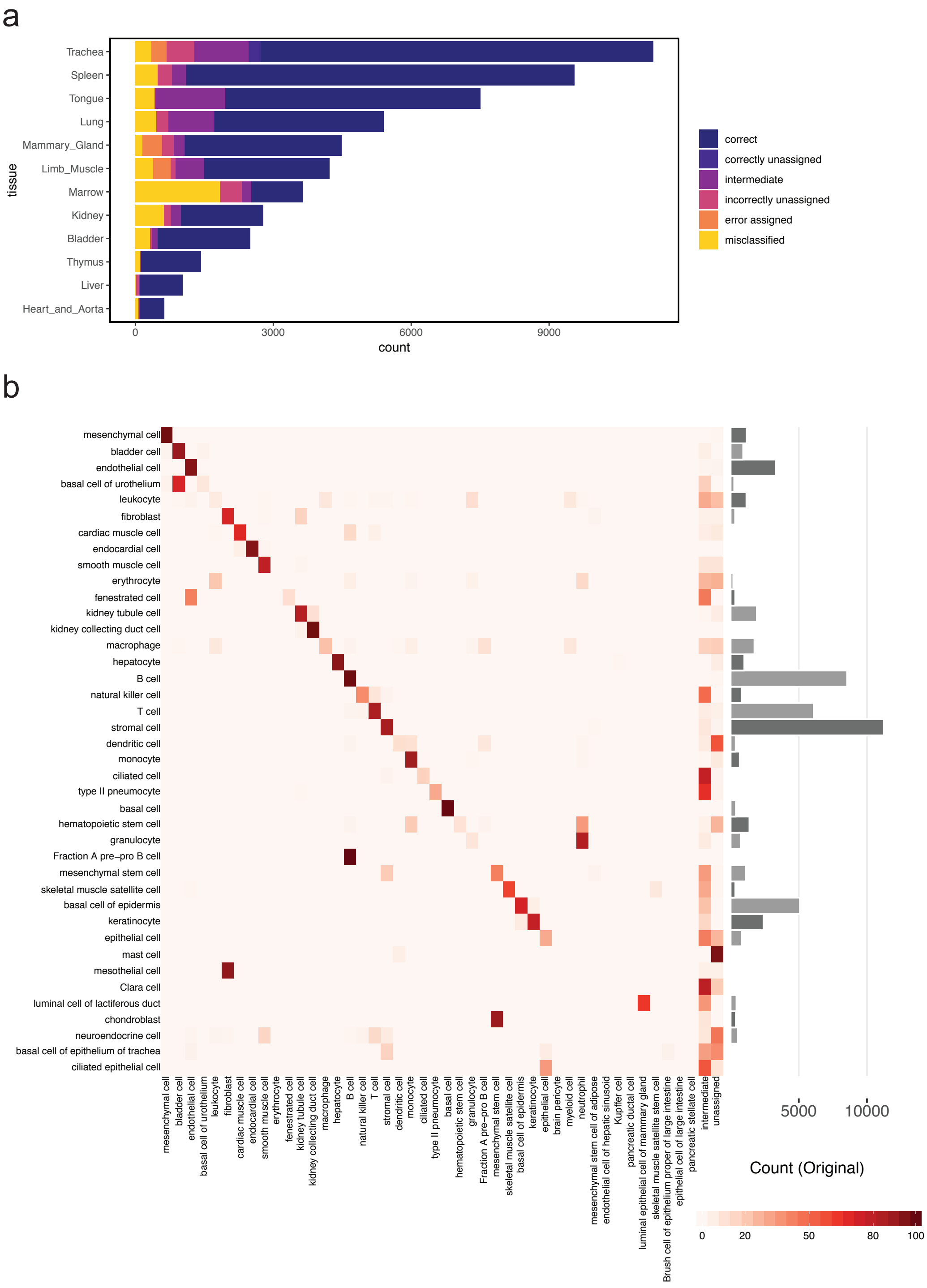


Supplementary Figure 6: (a) Barplot indicating the predicted cell types organized by tissue type when the Tabula Muris Microfluidic dataset is used as query and the Tabula Muris FACS data set is used as reference. (b) Heatmap of data in Fig. 2c comparing the original cell types given in the Tabula Muris Microfluidic data (rows) against the scClassify predicted cell types (columns) generated using the Tabula Muris FACS data as the reference dataset.

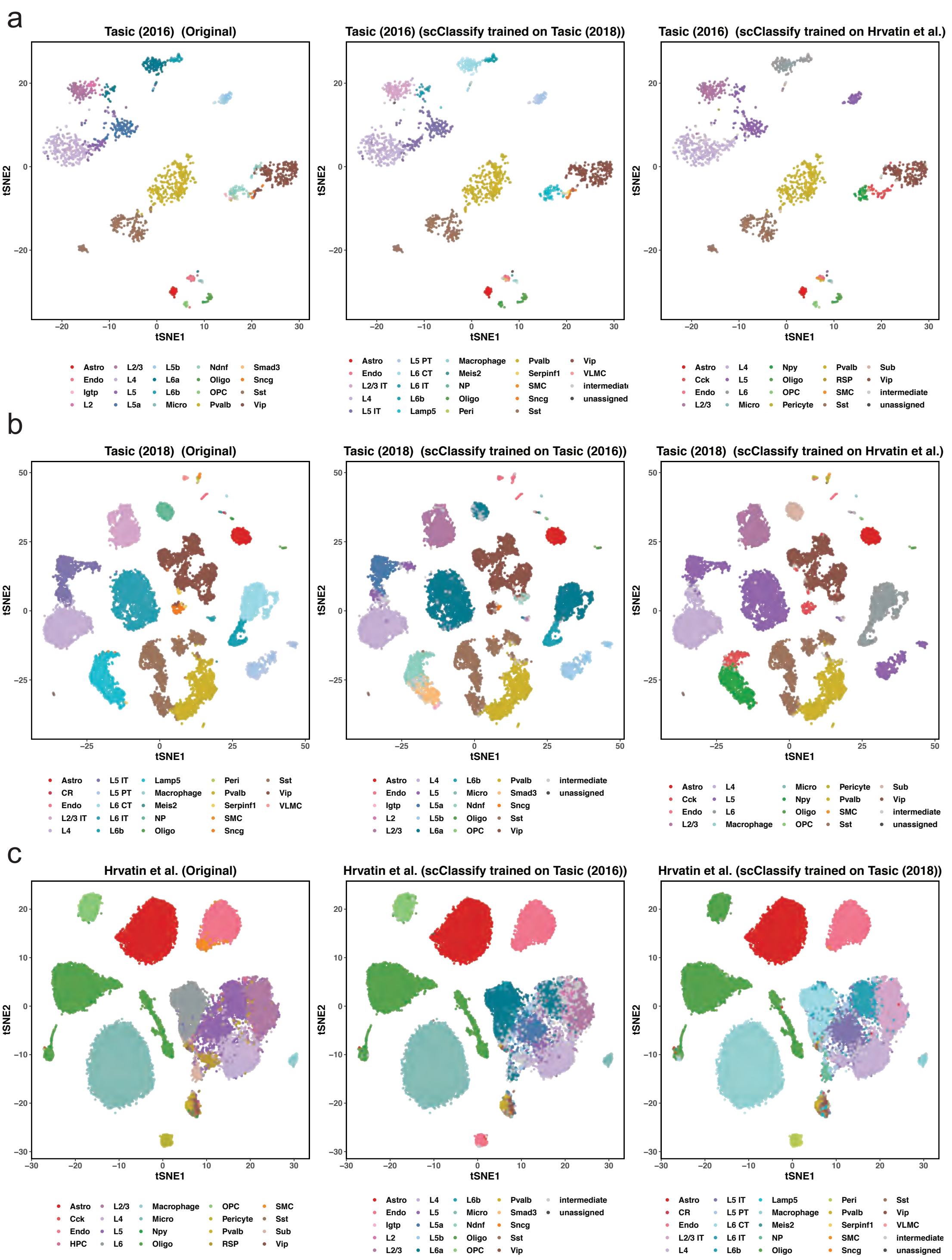


Supplementary Figure 7: (a) A 1 by 3 panel of tSNE plots of the Tasic et al. dataset (2016) from the neuronal data collection, where data points are color coded by original cell types given in [4] (left panel), the scClassify predicted cell types generated using Tasic et al. (2018) as the reference dataset (middle panel) and the scClassify predicted cell types generated using Hrvatin et al. as the reference dataset (right panel). (b) A 1 by 3 panel of tSNE plots of Tasic et al. (2018) from the neuronal data collection color coded by the original cell types given in [5] (left panel), the scClassify predicted cell types generated using Tasic et al. (2016) as the reference dataset (middle panel) and the scClassify predicted cell types generated using Hrvatin et al. as the reference dataset (right panel). (c) A 1 by 3 panel of tSNE plots of Hrvatin et al. from the neuronal data collection color coded by the original label [2] (left panel), the scClassify predicted cell types generated using Tasic et al. (2016) as the reference dataset (middle panel) and the scClassify predicted cell types generated using Tasic et al. (2018) as the reference dataset (right panel).

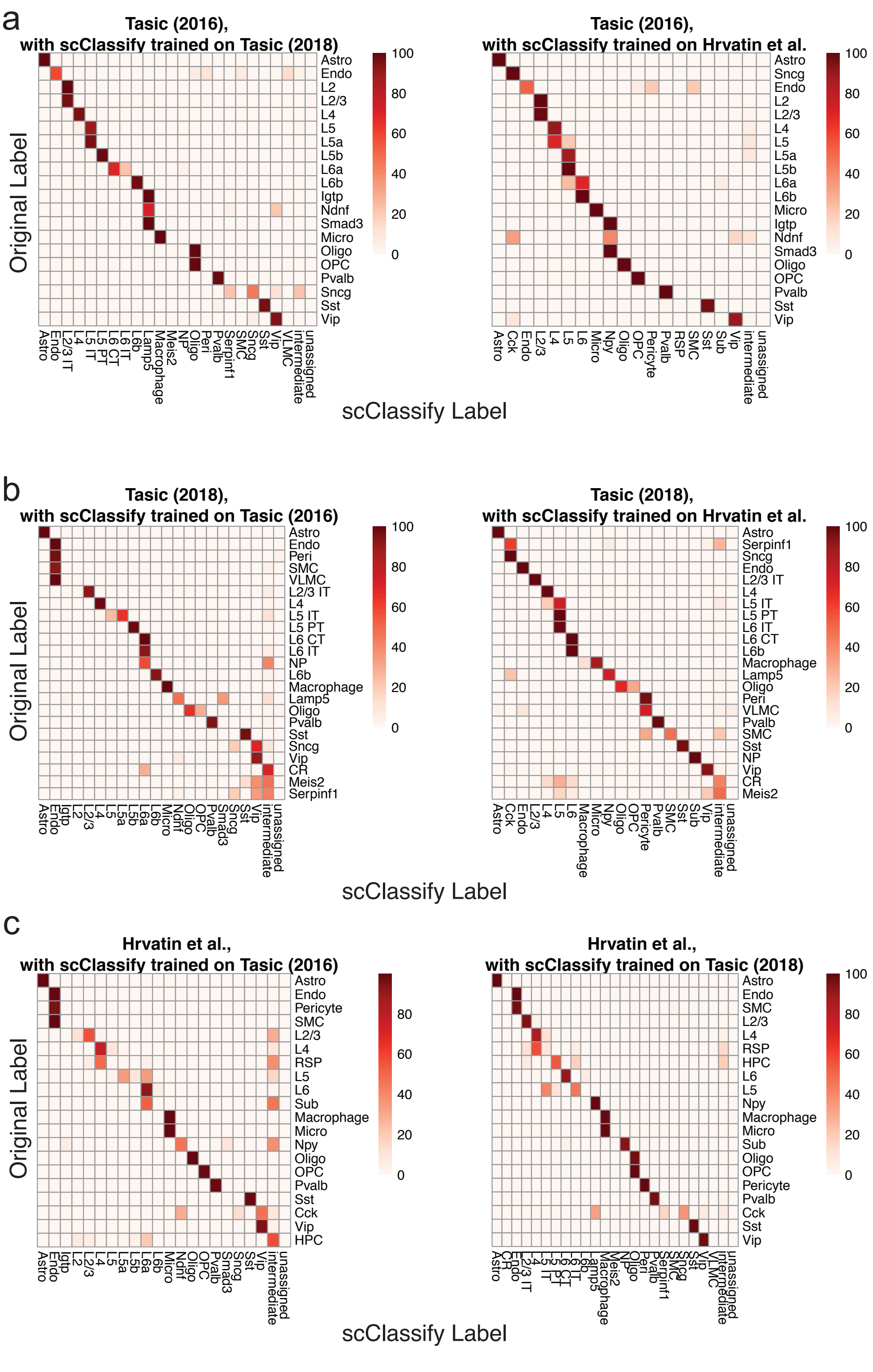


Supplementary Figure 8: (a) A 1 by 2 panel heatmaps of data in Supplementary Figure 6a comparing the cell types from the original cell types given in Tasic (2016) (rows) against scClassify predicted cell types (columns) generated using either the Tasic (2018) (left panel) or the Hrvatin et al. (right panel) as reference dataset. The squares are colored by the percentage of cells of a certain Tasic (2016) cell type. (b-c) as above for Supplementary Figures 7b and 7c.

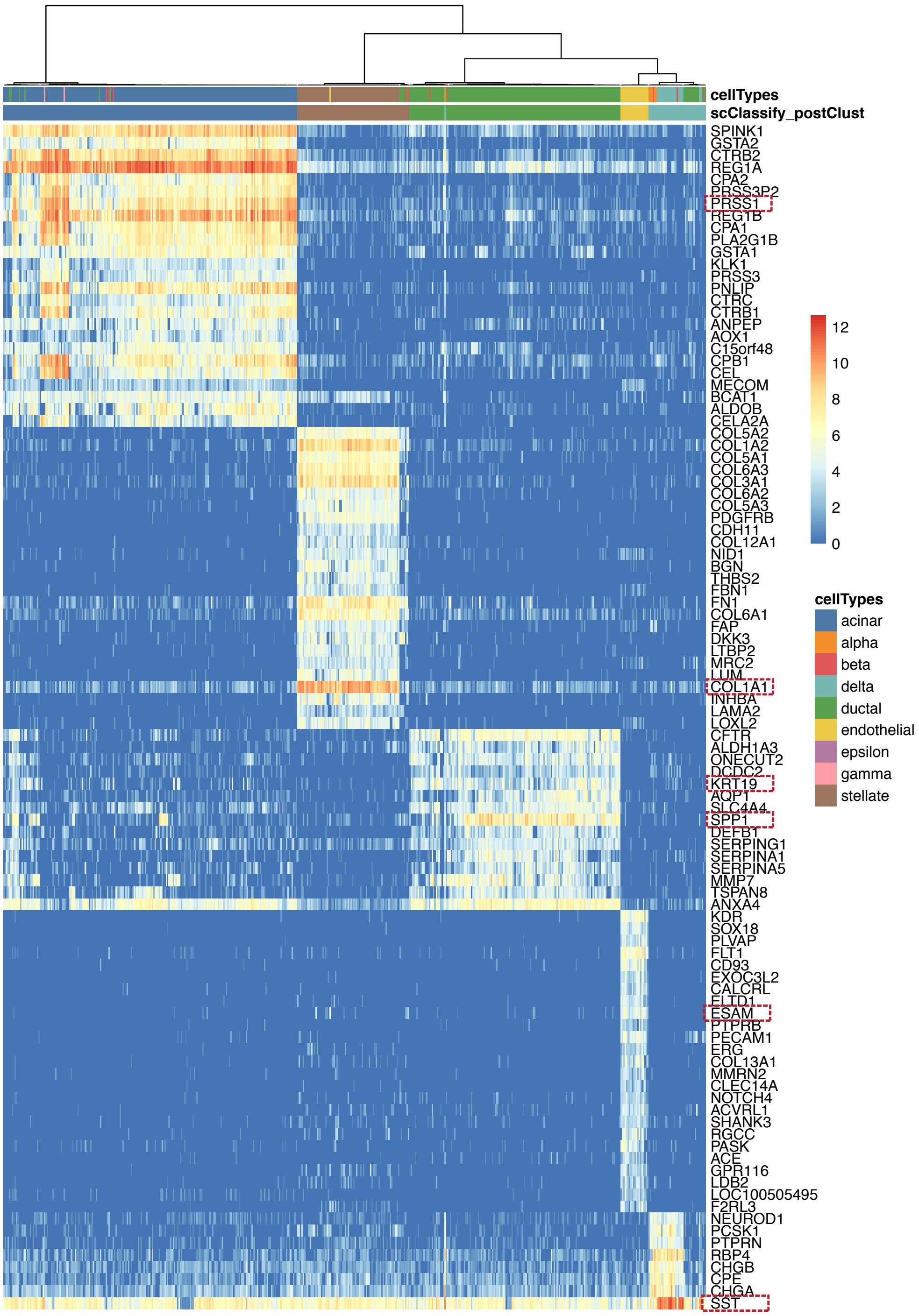


Supplementary Figure 9: Heatmap of the top 20 differentially expressed genes from each of the five cell-type clusters generated through post-hoc clustering of the Xin-Muraro data pair. Here, Xin et al. data is used as the reference dataset and Muraro et al. data as the query dataset. The heatmap is colored by the log-transformed expression values. The red rectangles indicate markers that are consistent with those found in the original study.

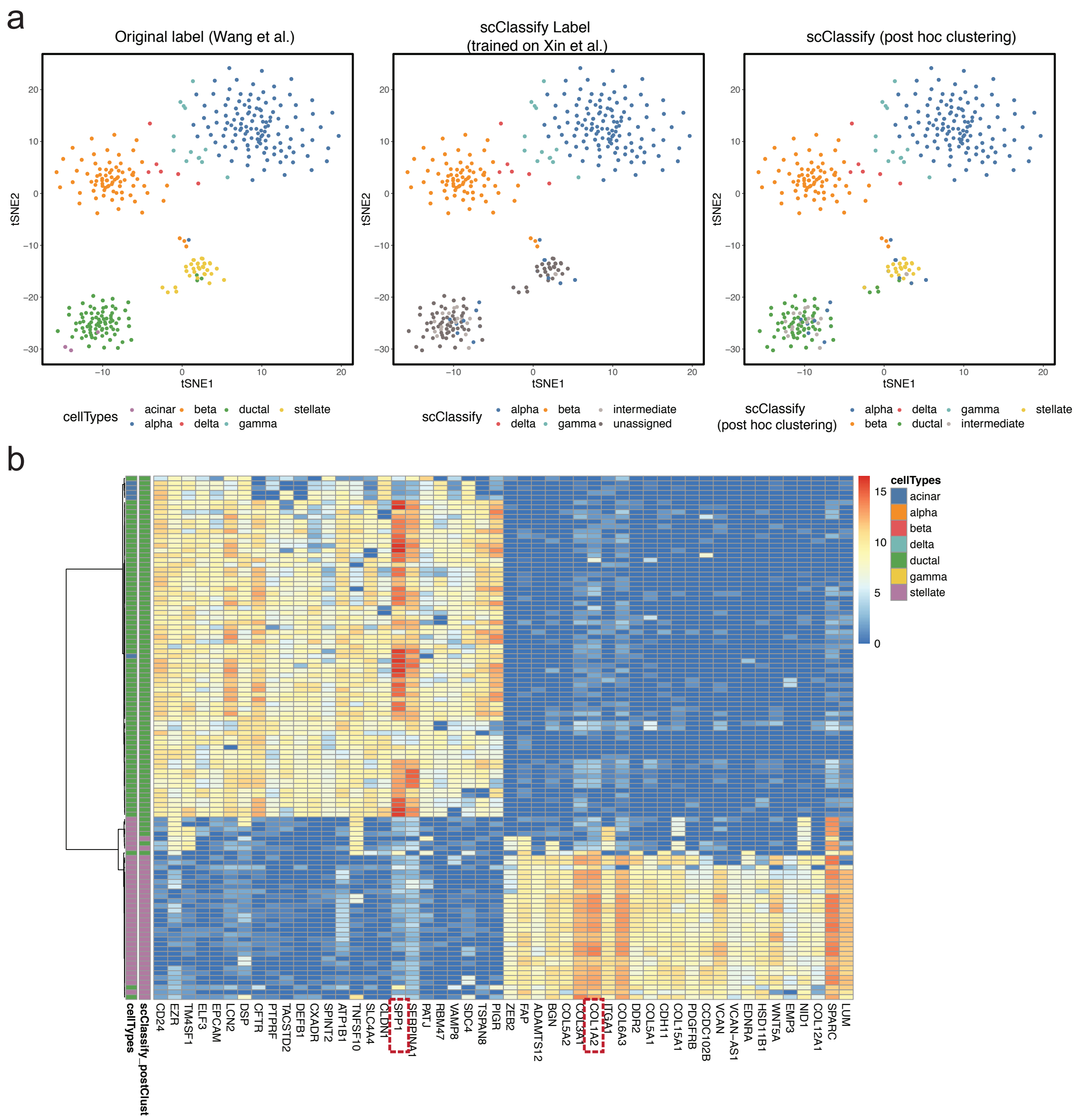


Supplementary Figure 10: (a) A 1 by 3 panel of tSNE plots of Wang et al. from the human pancreas data collection color coded by original cell types given in [7] (left panel), the scClassify label generated using Xin et al. as the reference dataset (middle panel) and the scClassify predicted cell types after performing post-hoc clustering (right panel). (b) Heatmap of the top 20 differentially expressed genes from each of the two cell-type clusters generated from post-hoc clustering of the Xin-Wang data pair. The heatmap is color coded by the log-transformed expression level. The red rectangles indicate markers that are consistent with those found in the original study.

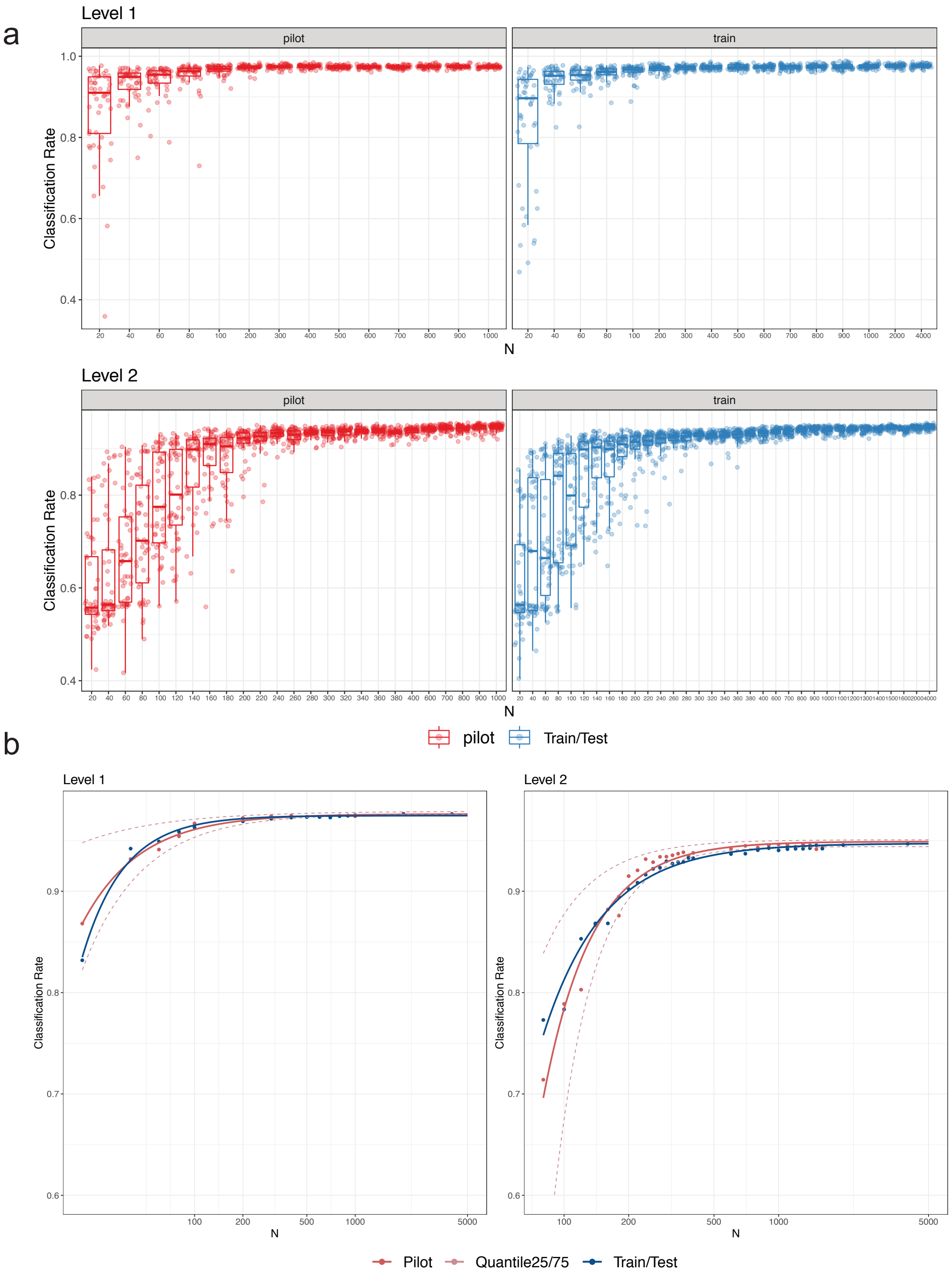


Supplementary Figure 11: (a) A 2 by 2 panel of collections of boxplots demonstrating the validation of the sample size calculation using the PBMC10k dataset. The x-axis indicates the sample size (N), and the y-axis indicates the accuracy rate. The left panel indicates the results for the pilot data (20% of the full dataset), and the right panel indicates the results for the reference-test data (the remaining 80% data), representing the data that would be obtained in a follow-up experiment. The top panel indicates the results of predicting PBMC at the top level of the cell type tree, while the bottom panel indicates the results of cell-type prediction at the second level of the cell type tree. (b) The fitted learning curves on the same data where red solid lines indicate the learning curves by fitting mean accuracy rate of pilot data; red dashed lines are the learning curves obtained by fitting the learning curves by fitting to the upper (75%) and lower (25%) quantiles of accuracy rate of pilot data. The blue lines indicate the learning curves by fitting the mean of the accuracy rate for the follow-up reference and test dataset.

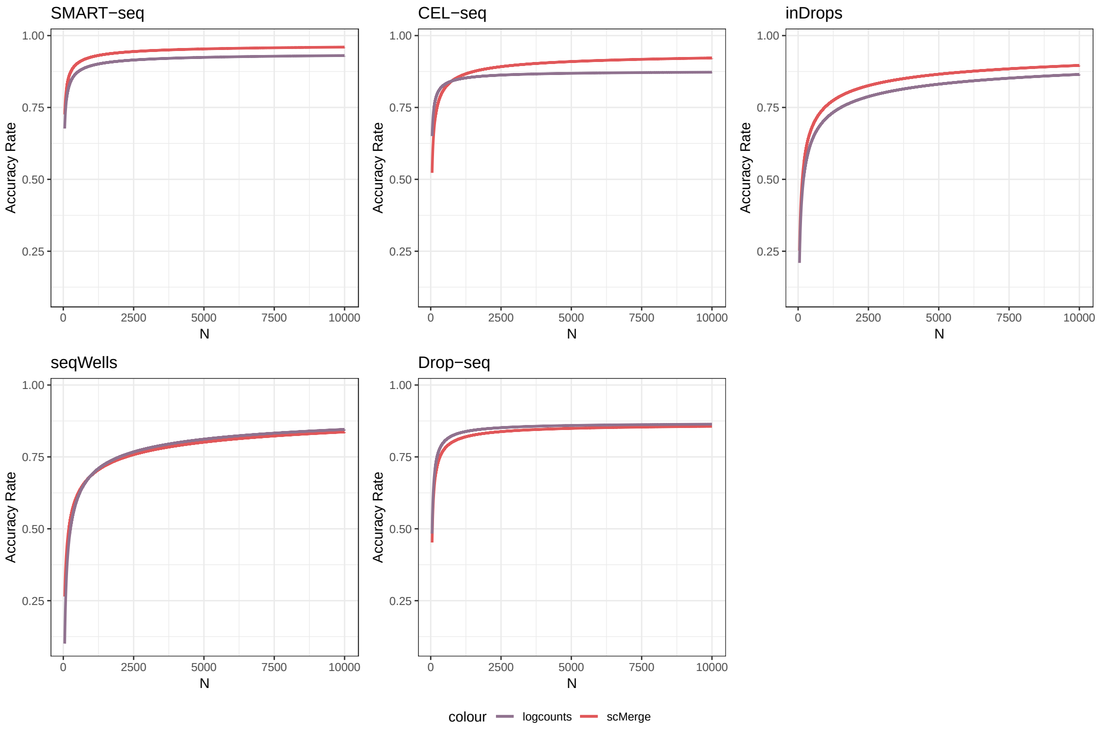


Supplementary Figure 12: Five learning curve plots illustrating the influence of batch effects on sample size requirement based on Ding et al. PBMC data generated from five different protocols: SMART-seq, CEL-seq, inDrops, seqWells and Drop-seq [1]. In each dataset, two or more PBMC samples displayed significant batch effect. Purple lines indicate the learning curves constructed using log-transformed data (with batch effect), and red lines denote the learning curve constructed after batch effect removal using scMerge [3].

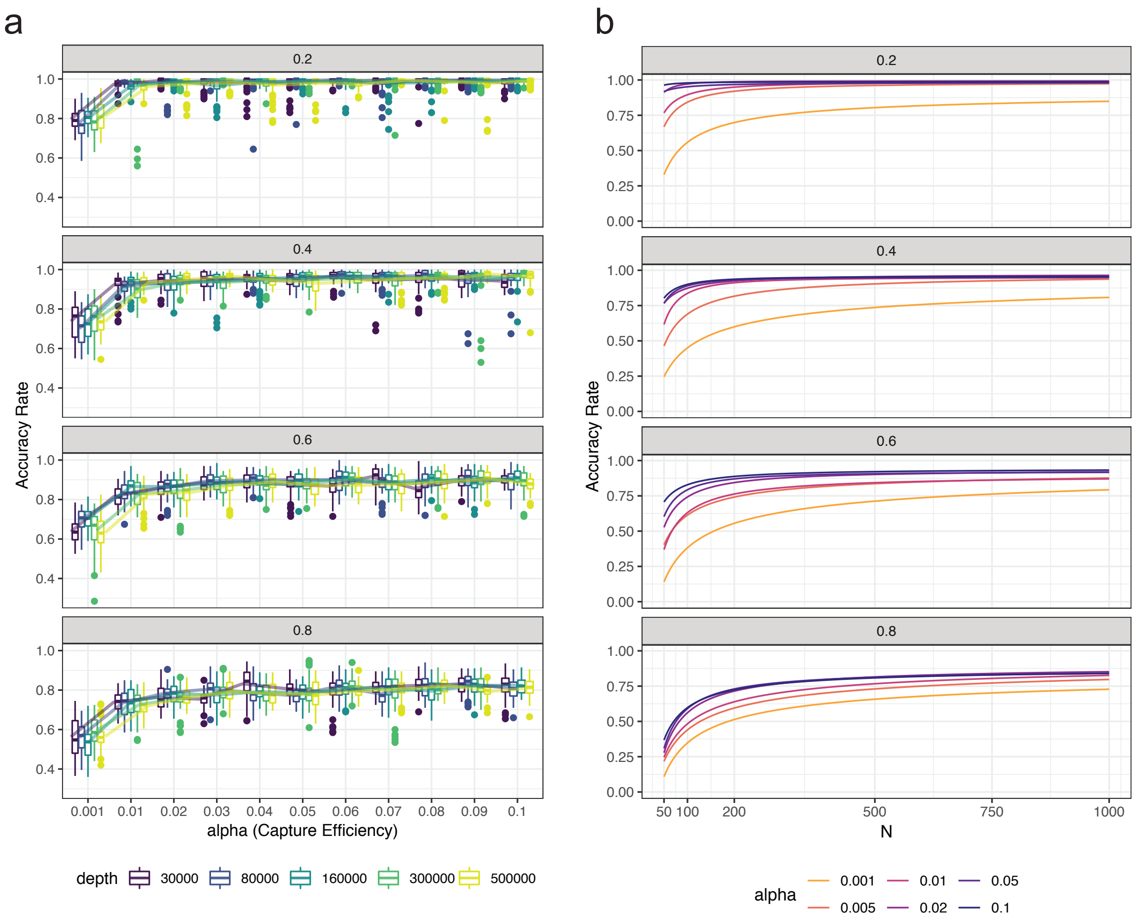


Supplementary Figure 13: (a) A 4 by 1 panel indicating the accuracy rate of the simulation results using SymSim [9] by estimating parameters from PBMC10k dataset. Each of four plots indicates accuracy rate with different degrees of within cell type heterogeneity (0.2, 0.4, 0.6, and 0.8), colored coded by five different sequencing depth (30000, 80000, 160000, 300000, 500000). The horizontal axis shows capture efficiencies ranged from 0.001 to 0.1, and y-axis indicates the accuracy rate. We found that the within-population heterogeneity has strong effect on the classification performance. scClassify can achieve above 95% average accuracy rate for the populations with high heterogeneity, while remaining at about 70% average accuracy rate for the extremely homogeneous population (sigma = 1). In terms of the capture efficiency, we found that the accuracy rates converge to values at about 0.02, even for extremely homogeneous population. (b) A 4 by 1 panel of fitted learning curves of the simulation results, where each plot indicates accuracy rate of different degrees of within cell type heterogeneity (0.2, 0.4, 0.6, and 0.8), colored coded by different capture efficiency (0.001, 0.005, 0.01, 0.2, 0.05, and 0.1). X-axis indicates the sample size (N) of the reference set, and y-axis indicates the accuracy rate. We found that for high heterogeneous population (sigma is 0.2, and 0.4) and a not so low capture efficiency rate (greater than 0.005), only small number of reference sample size is required to achieve a very high accuracy rate (around 95%). For a population that is more homogeneous, a higher capture efficiency rate will be required to achieve the greater accuracy rate that scClassify can achieved, as more reference samples are required since scClassify is learn relatively more slowly.

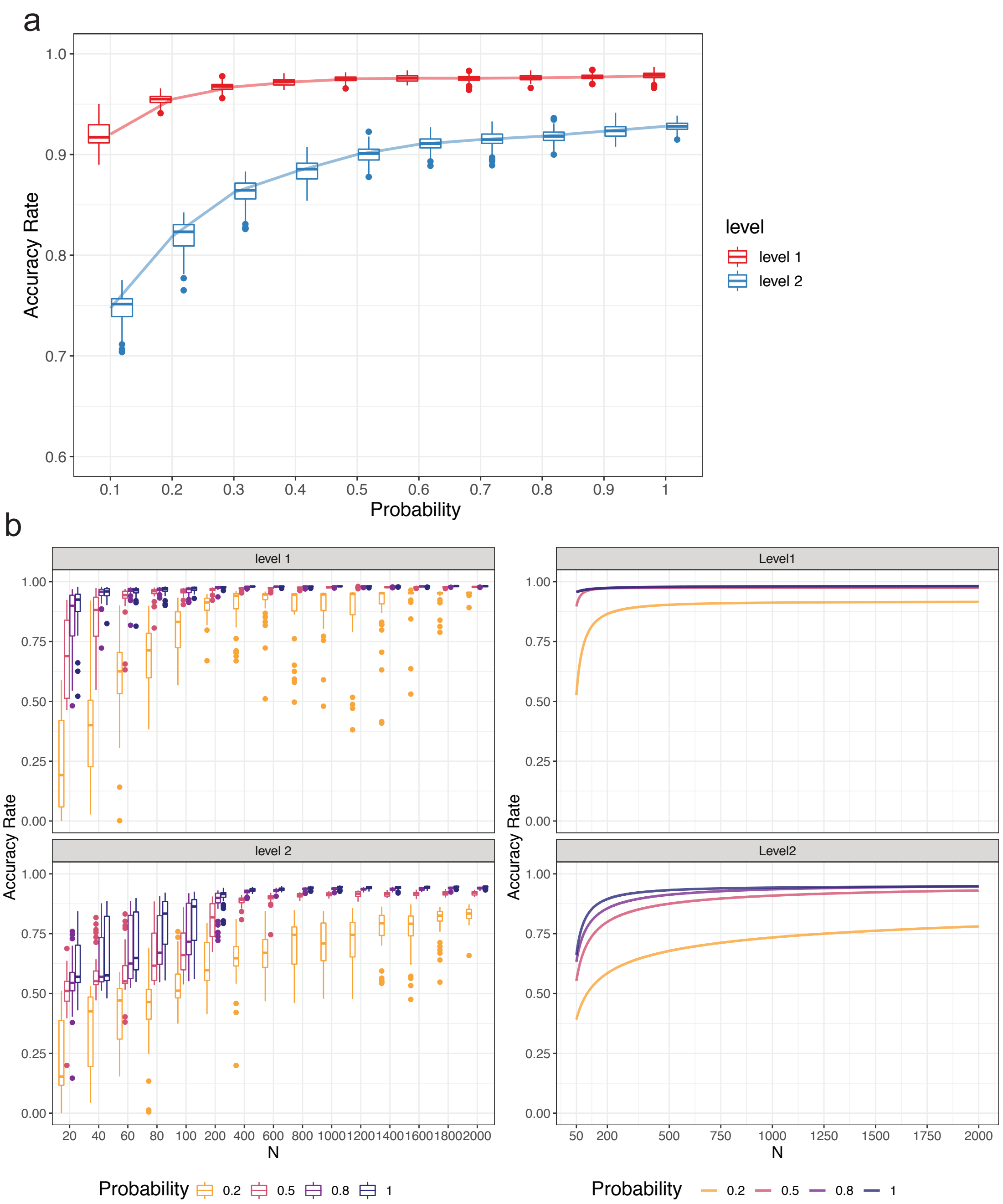


Supplementary Figure 14: Downsampling of the PBMC10k data using DECENT's beta-binomial capture model [8]. (a) Boxplots indicate the accuracy rates of the cell predictions from downsampled data for the top (red) and second level (blue) of the cell-type tree. The x-axis indicates the downsampling parameter of the beta-binomial distribution (that is, the ratio of capture efficiency in the downsampled dataset relative to the original dataset), and the y-axis denotes the accuracy rate. (b) Sample size calculation of down sampling. The left panel indicates the accuracy rate generated by repeating the training and testing procedure 50 times with varying size of the reference data and probabilities for down-sampling. The right panel displays the fitted learning curves based on the mean accuracy rate of the left panel. Both boxplots and lines are colored by probability of down-sampling (0.2, 0.5, 0.8 and 1). The top panel shows the results from the cell type predictions at the top level of the cell type tree, and the bottom panel shows the results from the cell type predictions at the second level of the cell type tree.

b

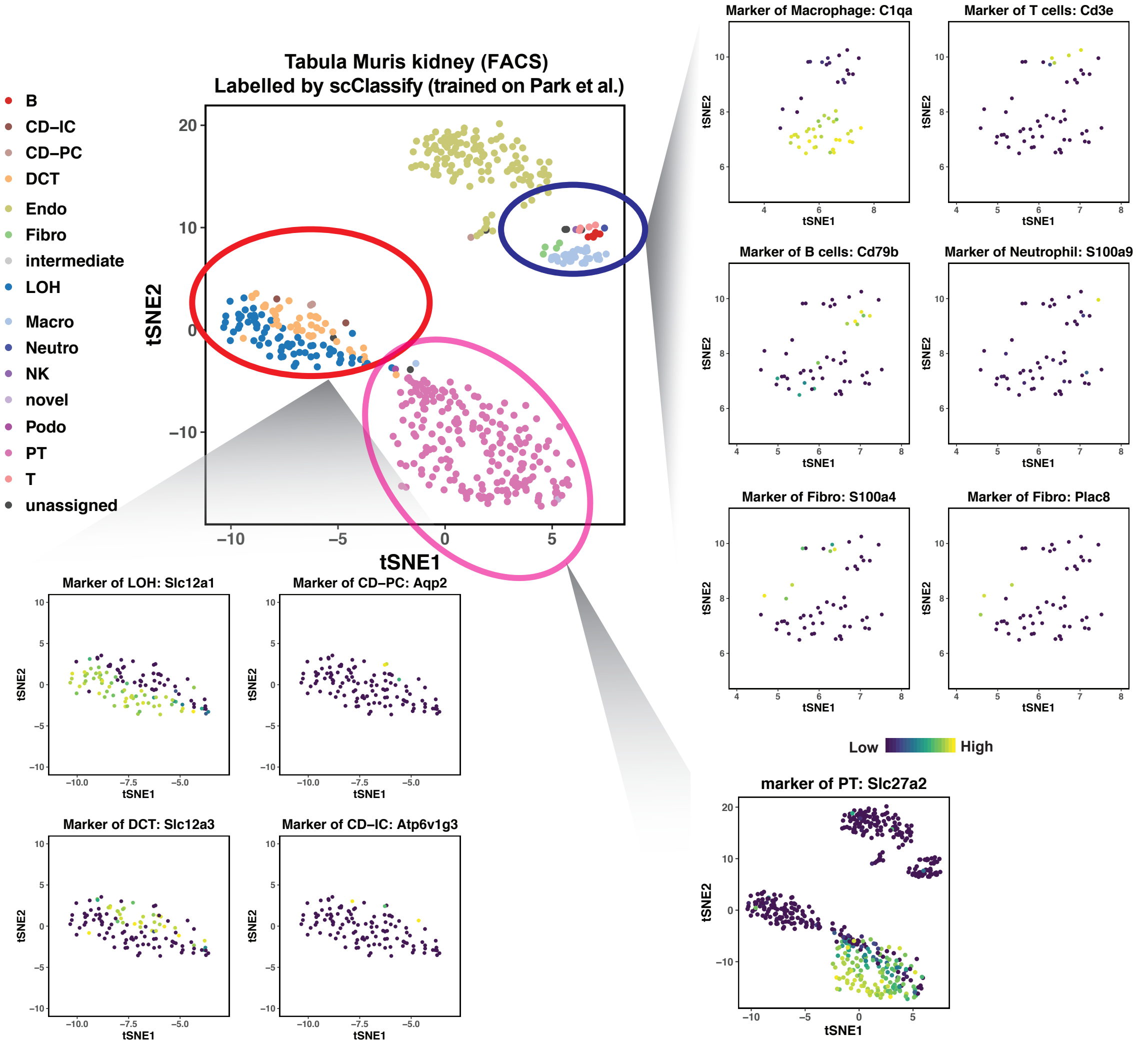


Supplementary Figure 15: scClassify results of the Tabula Muris Kidney FACS as query and the Park et al. kidney data as reference. The large tSNE plot displays the full data colored by the predicted cell types from scClassify, and the smaller marker expression plots are highlighted by the level of expression of 11 marker genes.

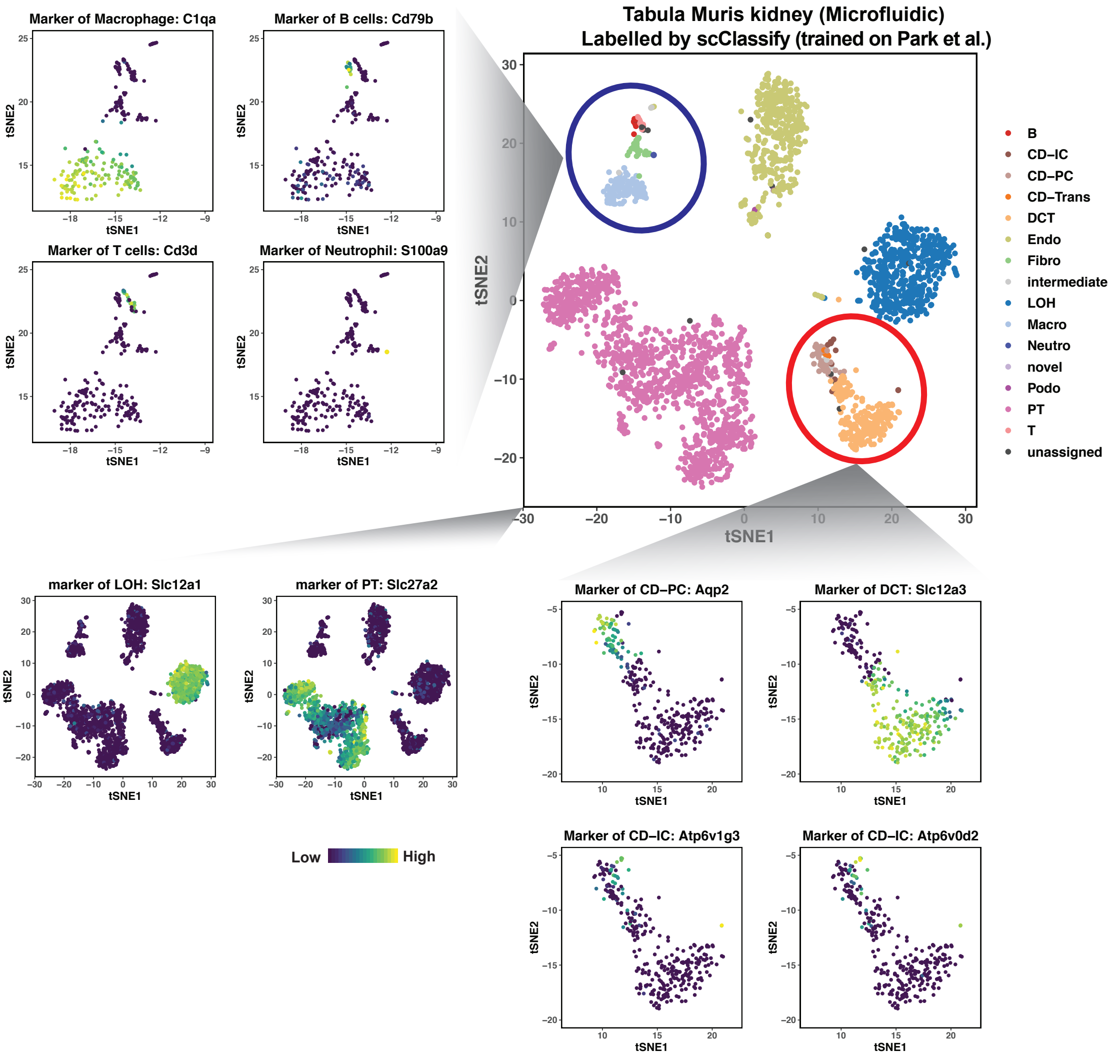


Supplementary Figure 16: scClassify results of the Tabula Muris Kidney Microfluidic data as query, and the Park et al. kidney data as reference. The large tSNE plot of the full data is colored by predicted cell types from scClassify, and the smaller marker expression plots are highlighted by the level of expression of 11 marker genes.
